# Rage bait: a neurofunctional signature of moral outrage elicited by social media

**DOI:** 10.64898/2026.11.18.767526

**Authors:** Xianyang Gan, Ran Zhang, Feng Zhou, Lisa Wagels, Heng Jiang, Junjie Wang, Yanan Qing, Xi Yang, Jingxian He, Zihao Zheng, Ting Xu, Klaus Mathiak, Bin Jiang, Dezhong Yao, Benjamin Becker

## Abstract

Moral outrage is ubiquitous online, with powerful emotional and motivational consequences for social life. Combining naturalistic fMRI during exposure to short-form social-media videos depicting moral violations with predictive modeling, we developed a neurofunctional signature of subjective moral outrage (MONS) in discovery/validation (*n*=80/30) cohorts. MONS predicted experienced outrage and punishment tendencies across cohorts. Positive predictive weights encompassed regions implicated in mentalizing, threat detection and action initiation (precuneus, posterior cingulate, periaqueductal gray, hypothalamus, cerebellum), whereas negative weights involved valuation and behavioral flexibility systems (amygdala, orbitofrontal cortex). Across 11 datasets (*n*=478), MONS captured anger and disgust but was dissociable from general negative affect and affective arousal. Notably, MONS did not predict perceived moral wrongness in conventional moral judgment paradigms, suggesting that moral outrage is not reducible to more cognitive moral evaluation. Findings provide a moral outrage neurofunctional model that delineates its relationship to other affective states and distinguishes it from traditional moral judgment.

## Introduction

Morality represents a central part of human sociality, from shaping social behavior and navigating relationships to mobilizing collective action, thereby serving as a cornerstone for the functioning and stability of human societies^1^. Beyond guiding judgments about right and wrong, morality is deeply intertwined with affective and motivational processes that are reflected in social emotions such as moral outrage^1^. Moral outrage, a mixture of anger and disgust^2–5^, is induced by stimuli appraised as violations of social norms^3,6^. The subjective experience of moral outrage (hereafter, “outrage”) motivates behavioral responses such as the punishment of moral transgressors^1,3,6–8^, which is vital for enforcing social norms^1,8–13^. Humans are evolutionarily predisposed to preferentially attend to moral and high-arousal negative stimuli because such information is motivationally relevant for social adaptation and biological survival^4,8,14–19^.

Much of human morality evolved in the context of small-group living^8^. Within these social environments, encounters with moral violations were relatively infrequent, and attention to such events served an adaptive function by enabling individuals to detect norm violations, experience outrage and engage in corrective actions such as punishment of wrongdoers^6,8,14^. However, social interactions are increasingly shifting into the online realm, with 5.81 billion people using social media as of October 2026^20^. The advent of the social media age has fundamentally transformed the context in which moral interactions take place relative to that of earlier human societies^1,8^. Unlike small-group environments in which moral information was largely constrained by direct social interactions and local communication networks, social media exposes individuals to a continuous stream of morally relevant information extending far beyond their immediate social circles. Consequently, an attentional system that evolved to prioritize rare but socially important moral violations is now operating within an environment characterized by an unprecedented abundance of moralized content^8^.

Social media is not merely a neutral communication platform; through its platform design features and recommendation algorithms, it systematically shapes human moral experiences and judgments^1,3–6,8,14,15,17,21–25^. This transformation is closely tied to the attention economy^26^, in which platforms compete for limited human attentional resources^15,27^ to maximize user engagement and, ultimately, generate revenue^1,14,15^. Because moral and high-arousal negative content is especially effective at capturing attention, recommendation algorithms exploit this preference by amplifying such content, including outrage-inducing material, thus sustaining engagement on platforms^1,4,6,8^. Indeed, converging evidence indicates that algorithms on major social media platforms (e.g., Meta, TikTok and Twitter/X) purposefully prioritize outrage content^28–30^. Consistent with this account, causal evidence demonstrates that engagement-based algorithms amplify moralized and emotional content, with the largest increases observed for outrage-related information^23^. Complementary causal evidence further shows that moral violations elicit outrage, which in turn increases preferences for social media engagement and facilitates viral spread^31,32^. These processes create a powerful self-reinforcing cycle^33^ where outrage-related content attracts attention, receives algorithmic amplification, and consequently gains increasing spread and visibility^3,14,17,34,35^. As a result, immoral acts are overrepresented on social media and evoke much stronger feelings of outrage than those individuals typically encounter in everyday offline life, thus disproportionately exposing people to extreme moral violations^1,4,6,8,17^. Beyond shaping outrage, social media may also intensify the punitive tendency toward transgressors^1,8,21^. The growing societal recognition of these dynamics is reflected in the selection of “***rage bait***” as the Oxford Word of the Year 2025^36^, underscoring increasing public awareness that outrage has become a powerful driver of attention, engagement and moral discourse in digital spaces.

Despite the growing societal importance of outrage in the social media age, little is known about its neurobiological basis. Existing accounts have largely relied on theoretical frameworks and large-scale analyses of online behavior, whereas the neural mechanisms through which moral-violation information elicits outrage and shapes subsequent punishment responses remain poorly understood. As humans become increasingly immersed in algorithmically curated digital environments, we continue to use brains tuned for offline small-group social living to navigate a novel informational ecosystem. Understanding how the brain represents outrage in contemporary digital contexts is therefore increasingly important. Recent advances in naturalistic neuroimaging paradigms, combined with multivariate pattern analysis (MVPA), provide a promising approach to address this question. Naturalistic neuroimaging enables the study of emotional responses elicited by ecologically relevant stimuli, whereas MVPA allows the characterization of their underlying neural representations. We capitalize on naturalistic functional magnetic resonance imaging (fMRI) using short-form videos depicting a wide range of moral transgressions collected from major short-form video platforms (e.g., Douyin, the Chinese counterpart of TikTok). Compared with the artificial moral-violation scenarios commonly used in studies of morality^37^, short-form videos are especially appropriate to ecologically elicit outrage in the scanner, as this form of content has become a dominant source of engagement and highly immersive information through user-friendly design and algorithm-promoted recommendations^38,39^. Such content often provokes outrage both online and offline. Meanwhile, by examining neural responses across multiple brain regions and voxels^40–43^, MVPA allows us to determine neural representations of mental processes with high resolution and considerably large effect sizes^42,44^. MVPA has successfully characterized neural representations of subjective emotional experiences^45,46^, including affective arousal and negative affect^47,48^.

Using naturalistic fMRI and MVPA, we addressed several key questions: (1) Is it possible to develop a neural signature sensitive to predict subjective outrage elicited by short-form videos with low to high outrage induction on the population level, and can this outrage signature further predict punishment tendencies (Fig. 1a,b)? (2) Can this signature predict trial-wise outrage experience on the individual level (Fig. 1c)? (3) Given that growing evidence from both human and animal research suggests that emotions are represented in distributed neural systems^45–49^, is subjective outrage similarly encoded in distributed brain systems or instead sufficiently captured by localized neural systems (Fig. 1d)? (4) Considering previous theoretical and behavioral accounts suggesting that outrage reflects a blend of anger and disgust^2–5,13,50,51^, is this relationship also evident on the neural level (Fig. 1e)? Next, we continued to test the discriminative validity of the developed moral outrage neural signature (MONS): (5) Prior conceptual and behavioral studies indicate conceptual overlap as well as separation between outrage and broader affective constructs such as negative affect and affective arousal^3,15,17^, is MONS distinct from established neural representations of negative affect^47^ and affective arousal^48^ or reducible to these more general affective representations (Fig. 1f)? (6) As the close behavioral links between outrage and moral judgment^35,50,52,53^, does MONS also encode perceived moral wrongness in classical moral judgment paradigms, which typically use written moral vignettes or static images as stimuli and emphasize the explicit cognitive evaluation of moral violations^54^ (Fig. 1g)? These questions provide a comprehensive framework for characterizing the neural representation of outrage and its role in punishment, affective processing and moral judgment in the social media age.

**Fig. 1.**
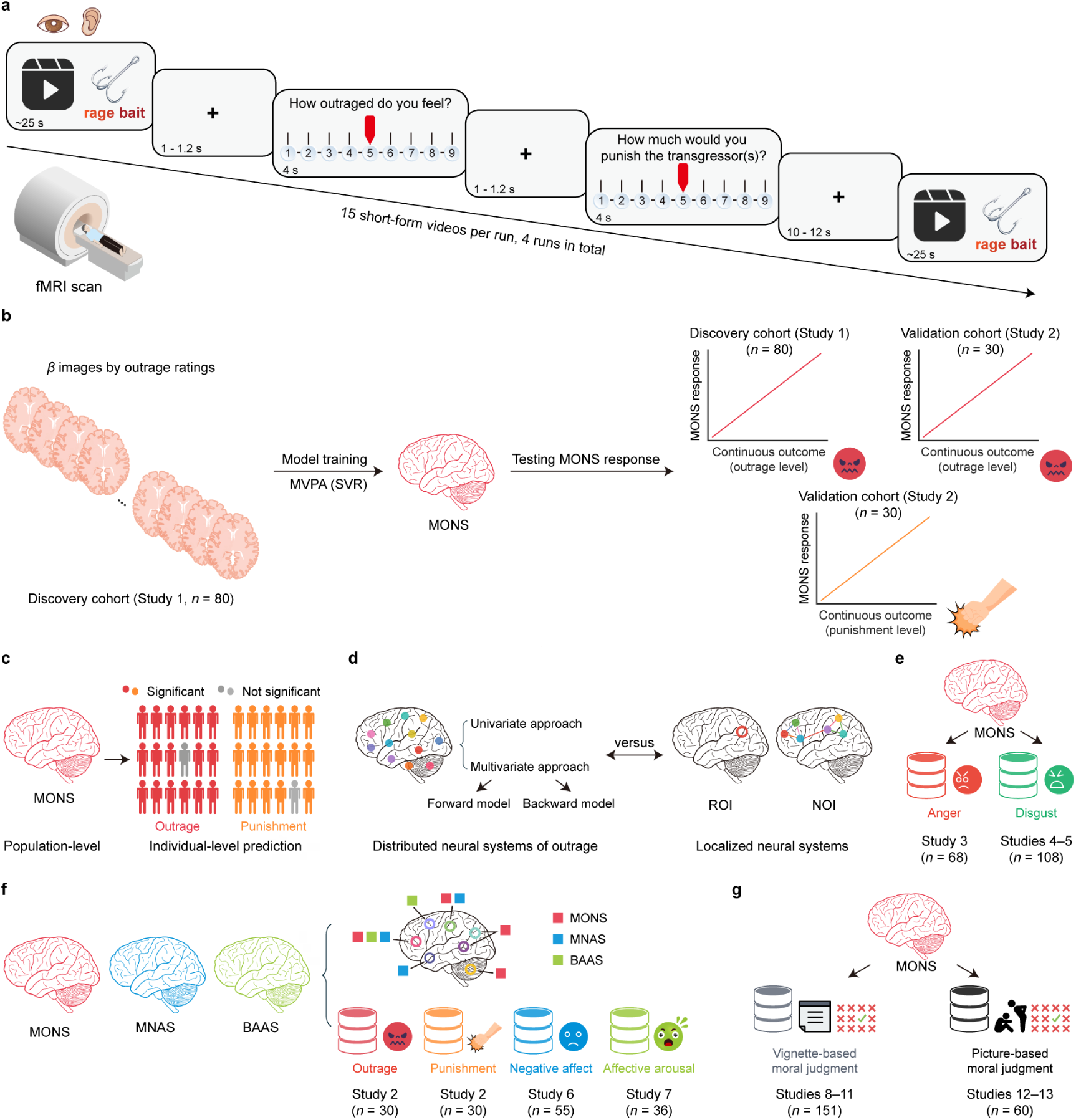
Task design and analytic workflow. **a**, Discovery and validation cohorts underwent the same naturalistic fMRI paradigm for outrage induction using short-form videos drawn primarily from Douyin and other social media platforms (Methods). **b**, Development and validation of the MONS. Voxel-level beta images were entered as features for model estimation. A whole-brain multivariate pattern predictive of the intensity of subjective outrage (MONS) was trained on the discovery sample (study 1, *n*=80) using SVR and evaluated using cross-validation in the discovery cohort. The trained model was then independently validated in a separate cohort (study 2, *n*=30) for its ability to predict both subjective outrage and punishment ratings. **c**, Within-subject trial-wise prediction using the population-level MONS to predict individual-level trial-wise outrage experience and punishment ratings in both discovery and validation cohorts. **d**, Characterization of the neural architecture underlying outrage using univariate and multivariate (forward/association and backward/prediction encoding^55^) approaches. Direct comparisons of distributed versus localized neural systems in predicting subjective outrage were then performed. **e**, Testing anger and disgust as components of outrage by applying the MONS to independent anger-related Taylor Aggression Paradigm (TAP)^56,57^ (study 3, *n*=68) and disgust^46^ (studies 4–5, *n*=108) datasets. **f**, Testing the discriminative validity of the MONS by conducting topographical and predictive comparisons with established signatures for negative affect (MNAS)^47^ and affective arousal (BAAS)^48^ (studies 6–7, *n*=55 and 36, respectively)^47,48^. **g**, Testing the discriminative validity of the MONS by applying it to perceived moral wrongness across six independent moral judgment datasets in artificial and static contexts^54^ (including four vignette-based datasets: studies 8–11, *n*=151; and two picture-based datasets: studies 12–13, *n*=60). BAAS=brain affective arousal signature from Zhang et al.^48^; MNAS=multimodal negative affect signature from Čeko et al.^47^; NOI=network of interest; ROI=region of interest; SVR=support vector regression.

## Results

### Behavioral results

We first examined whether the naturalistic audiovisual stimuli evoked varying levels of the subjective experience of outrage. During fMRI, participants viewed validated outrage-inducing short-form videos collected from major social media platforms and provided trial-wise ratings of outrage and punishment toward the moral transgressor(s) (if any) on 9-point Likert scales (1=neutral/slightest outrage or no punishment; 9=strongest outrage or severest punishment; see Methods). Ratings for both outrage and punishment were predominantly skewed toward the higher end of the scale (i.e., 6–9), with relatively fewer observations in the lower range (2–5), consistent with the intended selection of highly outrage-evoking stimuli (Supplementary Table 1). To ensure a sufficient number of trials per condition for subsequent analyses, ratings were grouped into five levels: 1–3 (level 1), 4–5 (level 2), 6–7 (level 3), 8 (level 4) and 9 (level 5). Detailed distributions of binned outrage and punishment ratings across cohorts are reported in the Supplementary Results (to account for potential effects of binning, we reproduced all analyses with the original ratings and results remained stable).

We next examined the relationship between subjective outrage and punishment tendencies. Outrage and punishment ratings were strongly correlated across the 45 outrage-inducing stimuli in both cohorts (discovery cohort: *r*=0.68±0.10; validation cohort: *r*=0.78±0.08), with correlations computed across participants for each stimulus and subsequently averaged across stimuli. These estimates were comparable to the association observed during the pre-rating procedure (*r*=0.68±0.23; Supplementary Table 2).

To further examine the relationship between outrage and punishment on the trial level, we constructed aligned rating profiles by sorting trials according to outrage ratings within each participant (Supplementary Methods). This procedure revealed a high degree of correspondence between the two measures across both cohorts. As outrage ratings increased monotonically across sorted trials, punishment ratings exhibited a closely matching trajectory, with substantial overlap between the two curves throughout the full range of ratings (Supplementary Fig. 1a,b). Together with the high within-subject correlations between the two measures observed in both cohorts (Supplementary Fig. 1c,d), these findings demonstrate a strong correspondence between trial-by-trial variations in experienced outrage and punishment tendencies.

### A brain signature for subjective experience of outrage (MONS)

We developed a whole-brain signature that sensitively predicts the subjective experience of outrage. In line with previous studies developing neuroaffective signatures^58,59^, we applied a linear SVR algorithm to identify a whole-brain signature of fMRI activity that tracked the intensity of self-reported outrage experience (Fig. 2a). The decoder was trained in the discovery cohort (*n*=80), next performance of the MONS was evaluated using 10×10-fold cross-validation within the discovery cohort and by determining reactivity of the MONS to outrage experience (as well as punishment tendency) in an independent validation cohort (*n*=30) that completed the same experimental paradigm.

**Fig. 2.**
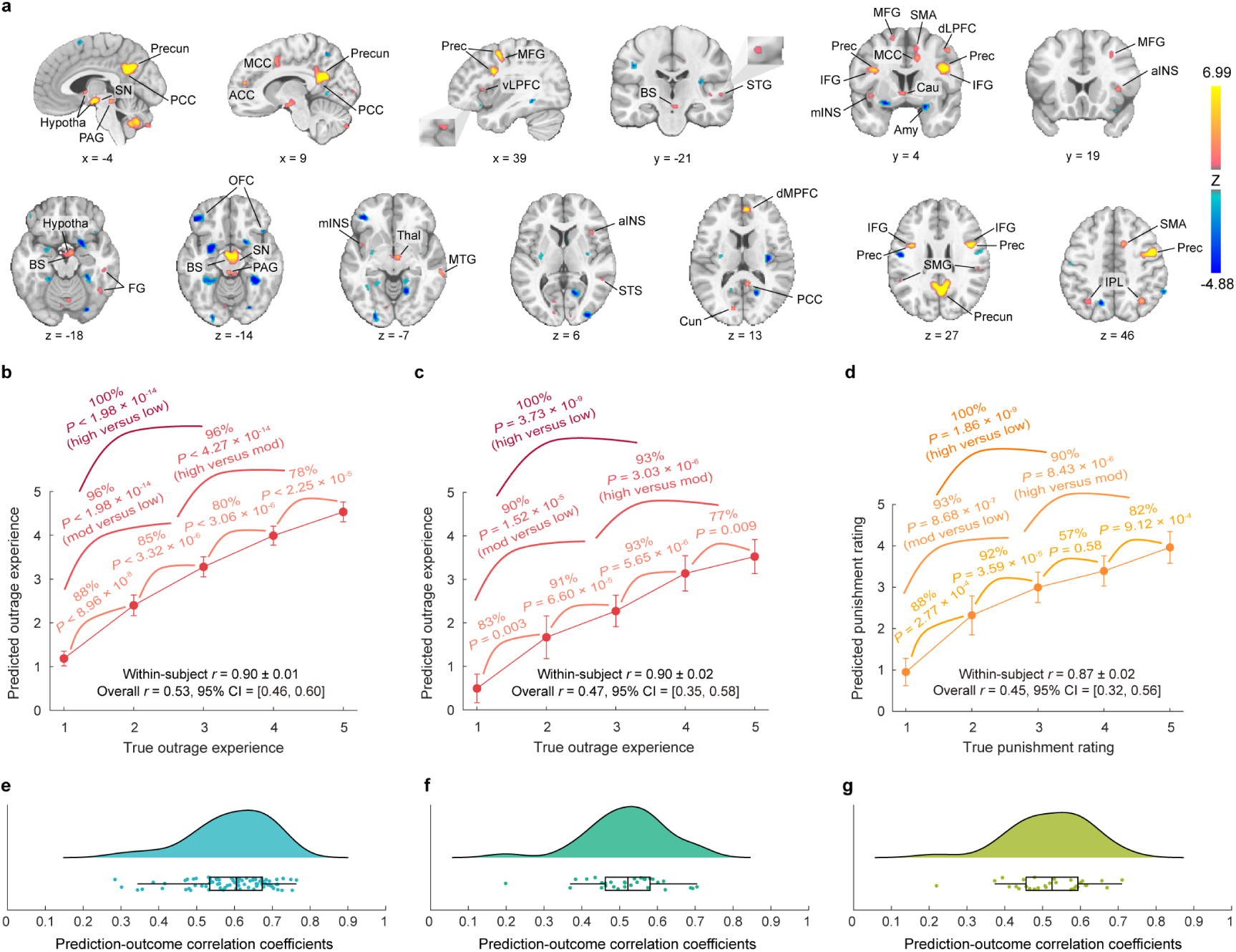
MONS. **a**, The MONS decoder thresholded at *q*<0.05 (FDR corrected, 10,000-sample bootstrap). **b**,**c**, Predicted outrage experience (subjective ratings; mean±s.e.m.) compared to actual outrage ratings in the cross-validated discovery cohort (**b**; *n*=80) and the independent validation cohort (**c**; *n*=30), respectively. **d**, Predicted punishment ratings (mean±s.e.m.) compared to actual punishment ratings in the independent validation cohort (*n*=30). **b**–**d**, Accuracy provided for forced-choice comparisons for different rating levels. *r* indicates the Pearson correlation coefficient between predicted and true ratings. **e**, Distribution of within-subject trial-wise Pearson correlations between predicted and true outrage ratings across participants in the discovery cohort, significant prediction–outcome correlations were observed in 79 of 80 participants (*P*=[1.28×10^−12^, 0.048]; remaining participant: *P*>0.05). **f**, Distribution of within-subject trial-wise Pearson correlations between predicted and true outrage ratings across participants in the validation cohort, significant prediction–outcome correlations were observed in 29 of 30 participants (*P*=[9.44×10^−10^, 0.004]; remaining participant: *P*=0.14). **g**, Distribution of within-subject trial-wise Pearson correlation between predicted and true punishment ratings across participants in the validation cohort, significant prediction–outcome correlations were observed in 29 of 30 participants (*P*=[6.09×10^−10^, 0.004]; remaining participant: *P*=0.10). **e**–**g**, In the raincloud plots, boxplots indicate the interquartile range (IQR), with the center line representing the median, box boundaries indicating the 25th and 75th percentiles, and the whiskers extending to the minimum and maximum values within 1.5 × IQR from the quartiles. The *P* values are based on two-sided independent binomial tests. Error bars indicate the s.e.m. ACC=anterior cingulate cortex; aINS=anterior insula; Amy=amygdala; BS=brainstem; Cau=caudate; Cun=cuneus; dLPFC=dorsolateral prefrontal cortex; dMPFC=dorsomedial prefrontal cortex; FG=fusiform gyrus; Hypotha=hypothalamus; IFG=inferior frontal gyrus; IPL=inferior parietal lobule; MCC=middle cingulate cortex; MFG=middle frontal gyrus; mINS=middle insula; MTG=middle temporal gyrus; OFC=orbitofrontal cortex; PAG=periaqueductal gray; PCC=posterior cingulate cortex; Prec=precentral gyrus; Precun=precuneus; SMA=supplementary motor area; SMG=supramarginal gyrus; SN=substantia nigra; STG=superior temporal gyrus; STS=superior temporal sulcus; Thal = thalamus; vLPFC=ventrolateral prefrontal cortex.

The MONS robustly predicted the level of subjective outrage in the discovery cohort (averaged within-subject correlation was 0.90±0.01 (mean±s.e.m.), average root mean squared error (RMSE) of 1.69±0.10; overall (between- and within-subject) prediction–outcome correlation was 0.53, averaged across ten repetitions, bootstrapped 95% confidence interval (CI) was [0.46, 0.60], Fig. 2b). Applying the predictive model trained on the discovery cohort–without further model fitting– to the validation cohort (using outrage ratings as the target variable) yielded similarly high predictive performance (within-subject *r*=0.90±0.02, average RMSE=1.92±0.20; overall *r*=0.47, 95% CI=[0.35, 0.58]; Fig. 2c), indicating a sensitive and robust neurofunctional signature of subjective outrage. When applied to the validation cohort with punishment ratings as the outcome measure, the MONS also demonstrated strong predictive performance (within-subject *r*=0.87±0.02, average RMSE=1.88±0.16; overall *r*=0.45, 95% CI=[0.32, 0.56]; Fig. 2d), suggesting that the MONS also captures punishment tendencies.

To further evaluate the MONS’s sensitivity to gradations in subjective outrage experience and punishment tendencies, we conducted within-subject forced-choice classification analyses based on pairs of activation maps (Supplementary Results). The MONS reliably distinguished different levels of outrage and punishment ratings, including adjacent rating levels, with accuracies substantially above chance across cohorts. These findings further demonstrate that the MONS captures graded variations in subjective outrage experience and associated punishment tendencies.

Given that audiovisual stimuli were used in this study and that previous evidence has shown that auditory and visual cortices contribute to encoding emotional information in both modalities^47^, we conducted lesion-based prediction analyses^60^ by excluding these sensory cortices. The resulting signature achieved similar performance, suggesting that outrage prediction was not primarily driven by information encoded in auditory and visual areas (Supplementary Results and Supplementary Fig. 2).

### Within-subject trial-wise prediction

Given that subjective outrage experience fluctuates dynamically across individuals, we next examined whether the population-level MONS could track moment-to-moment variations in outrage experience on the individual trial level. Single-trial MONS expression values were correlated with corresponding outrage ratings within each participant (Methods). Statistical significance was assessed using two-sided Pearson prediction–outcome correlation. In the discovery cohort, the MONS significantly predicted trial-by-trial outrage ratings in 79 of 80 participants (cross-validated), yielding a mean prediction–outcome correlation of *r*=0.59±0.01 (mean±s.e.m.; Fig. 2e). Consistent effects were observed in the independent validation cohort, where the MONS significantly predicted outrage ratings in 29 of 30 participants (mean *r*=0.52±0.02 (Fig. 2f). The MONS also predicted trial-wise punishment ratings in 29 of 30 validation participants (mean *r*=0.52±0.02; Fig. 2g). Combined, these findings demonstrate that the MONS captures dynamic, trial-level fluctuations in subjective outrage experience and related punishment responses. Importantly, similar predictive performance was observed when using the original, ungrouped ratings (1–9) as trial-wise outcomes, providing further support for the rating-grouping approach adopted in the primary analyses (Supplementary Results and Supplementary Fig. 3).

### Regions associated with and predictive of subjective outrage

We next characterized the neural systems contributing to the whole-brain prediction of subjective outrage. To identify brain regions that reliably contributed to the MONS, we first performed a bootstrap test to determine regions with significant, stable model weights (*q*<0.05, FDR corrected)^60^. Because some brain features may contribute to controlling for noise in the data^55^ rather than the target emotional process per se, we next transformed the population-level MONS into ‘activation pattern’ (‘structure coefficient’, see Methods). Both analyses revealed a distributed set of cortical and subcortical regions exhibiting significant model weights (*q*<0.05, FDR corrected; Fig. 3a) and structure coefficients (*q*<0.05, FDR corrected; Fig. 3b), including subcortical regions implicated in affective and threat-related processing, such as the PAG, brainstem, hypothalamus and thalamus^9,61,62^; regions involved in social and self-referential cognition, encompassing the precuneus, PCC, MCC, dMPFC, temporoparietal junction (TPJ) and lateral temporal cortices (e.g., superior temporal sulcus, superior temporal gyrus, middle temporal gyrus, temporal pole)^9,62–67^; and frontal, insular, parietal and motor regions implicated in cognitive control, interoception and action preparation, comprising the ACC, inferior frontal gyrus, middle frontal gyrus, posterior insula, supplementary motor area, precentral gyrus, inferior parietal lobule, supramarginal gyrus, substantia nigra and cerebellar regions^9,62,68–74^; additional regions included the caudate, fusiform gyrus and cuneus (Fig. 3c). Brain regions associated with and predictive of subjective outrage on the individual level, identified using convergent univariate and multivariate approaches, revealed a highly similar set of broadly distributed regions (Supplementary Methods, Supplementary Results and Supplementary Fig. 4). Overall, these convergent analyses indicated that subjective outrage is represented in a broadly distributed neural architecture spanning affective, social-cognitive and action-related systems. Negative associations with subjective outrage were most consistently observed in the OFC and occipital lingual regions. Complementary analyses further showed that the whole-brain MONS outperformed models restricted to individual regions or large-scale networks, supporting a distributed rather than localized neural representation of subjective outrage (see Supplementary Results, Supplementary Fig. 5,6 and Supplementary Tables 3,4).

**Fig. 3.**
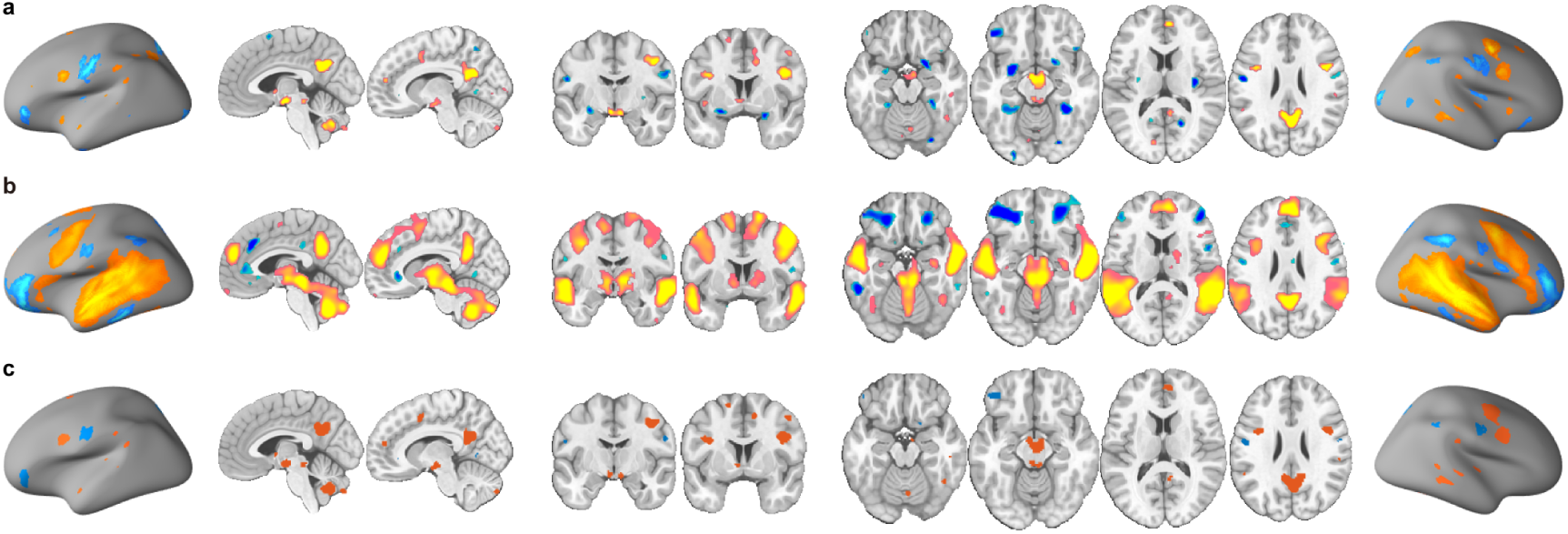
The subjective experience of outrage is associated with and predicted by distributed brain regions. **a**, Thresholded MONS. **b**, Thresholded transformed MONS ‘activation pattern’. **c**, Conjunction between MONS and transformed ‘activation pattern’. Images thresholded at *q*<0.05, FDR corrected. Hot color indicates positive predictive weights (**a**) or associations (**b**), whereas cold color indicates negative predictive weights (**a**) or associations (**b**).

### Evaluating the neurobiological validity of the signature

To further evaluate the neurobiological plausibility of the developed model, we calculated the spatial overlap percentage between the thresholded MONS and the modified 279-region version of the Brainnetome Atlas^75,76^ (see Methods). The top 20 atlas regions visualized in Fig. 4a included three precuneus sub-regions, two precentral gyrus sub-regions, two cingulate cortex sub-regions, two superior frontal gyrus sub-regions, two inferior parietal lobule sub-regions, fusiform gyrus, hypothalamus, brainstem, PAG and five cerebellar subregions. Functional decoding utilizing the BrainMap database confirmed that the identified brain systems were primarily associated with social cognition, negative affect (e.g., anger), arousal and salience (e.g., attention), general cognitive processing (e.g., cognition, memory, language, reasoning) and action responses (e.g., motion, motor learning, execution, inhibition) (Fig. 4b).

**Fig. 4.**
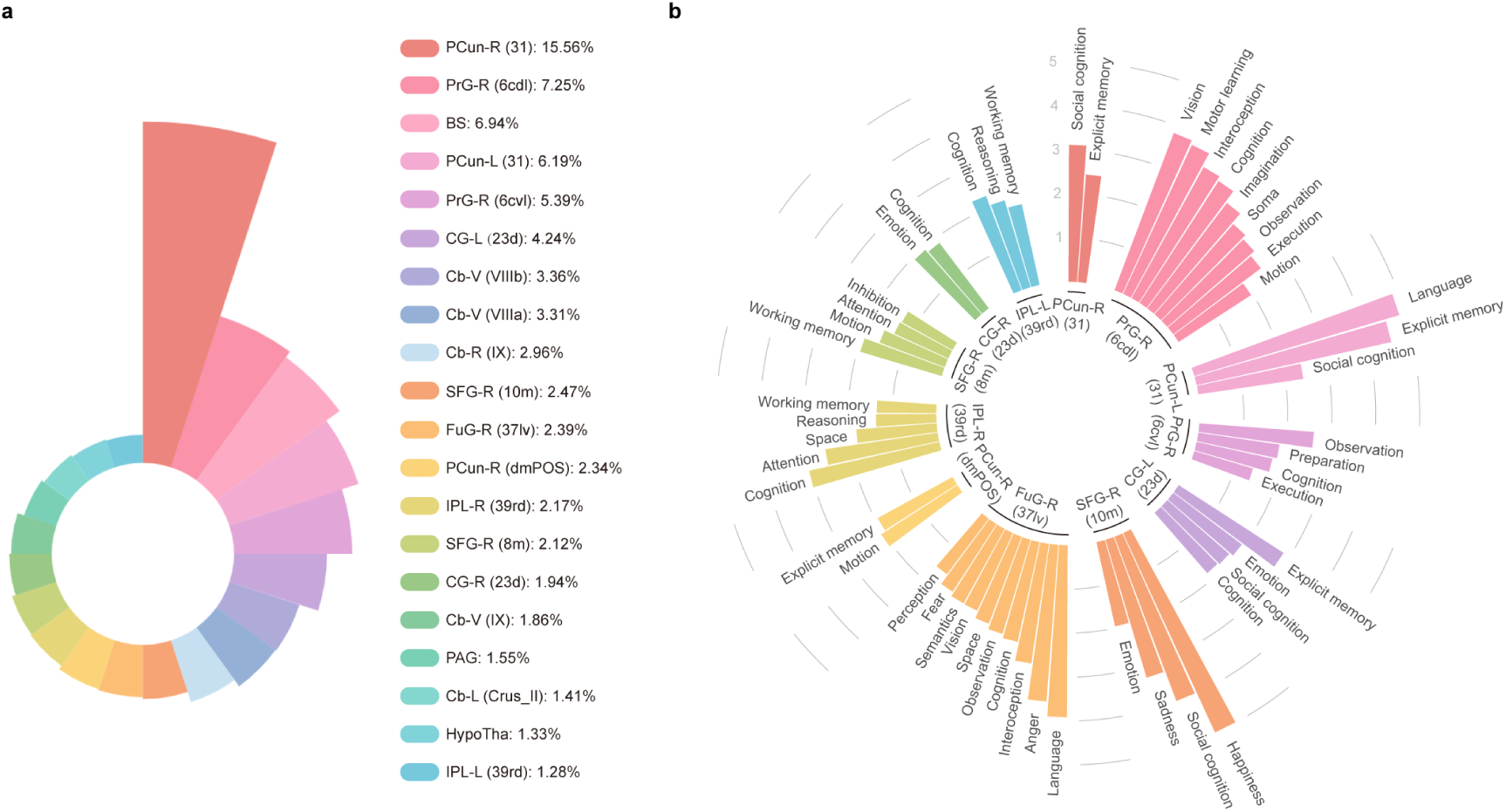
Neurobiological validity of the thresholded MONS. **a**, The top 20 atlas regions overlapping with the thresholded MONS (retaining positive values). **b**, Functional behavioral decoding for 12 exemplary regions for the thresholded MONS (provided by the Brainnetome website).

### MONS captures anger- and disgust-related components

Both theoretical accounts and empirical evidence^2–5,13,50,51^ suggest that outrage reflects a mixture of anger and disgust. We therefore tested whether the MONS was sensitive to neural responses associated with these two affective components. In the provocation-based TAP dataset^56,57^ (study 3, Supplementary Table 5), which assessed reactive aggression (retaliatory punishment) together with changes in state anger following the task (Methods), the MONS significantly differentiated aggression (money subtractions) following high versus low provocation among participants who reported increased post-task state anger (*n*=41, accuracy=71%, *P*=0.01, area under the curve (AUC)=0.71, Cohen’s *d*=0.52), but not among those without increased state anger (*n*=27, accuracy=44%, *P*=0.70, AUC=0.52, Cohen’s *d*=0.006; Fig. 5a). We next tested the MONS in two independent disgust datasets^46^ acquired while participants viewed a series of visual disgust stimuli eliciting different levels of subjective disgust experience (study 4, *n*=78; study 5, *n*=30; Supplementary Table 5). The MONS significantly discriminated different levels of subjective disgust in both datasets, with significant discrimination across most disgust-level contrasts (accuracy=[70%, 90%], Cohen’s *d*=[0.43, 1.17]) (Fig. 5b,c and Supplementary Table 6). Together, these findings indicate that the neural signature of outrage is sensitive to both anger-related aggression and subjective disgust, providing neurobiological evidence for the proposed anger–disgust components of outrage.

**Fig. 5.**
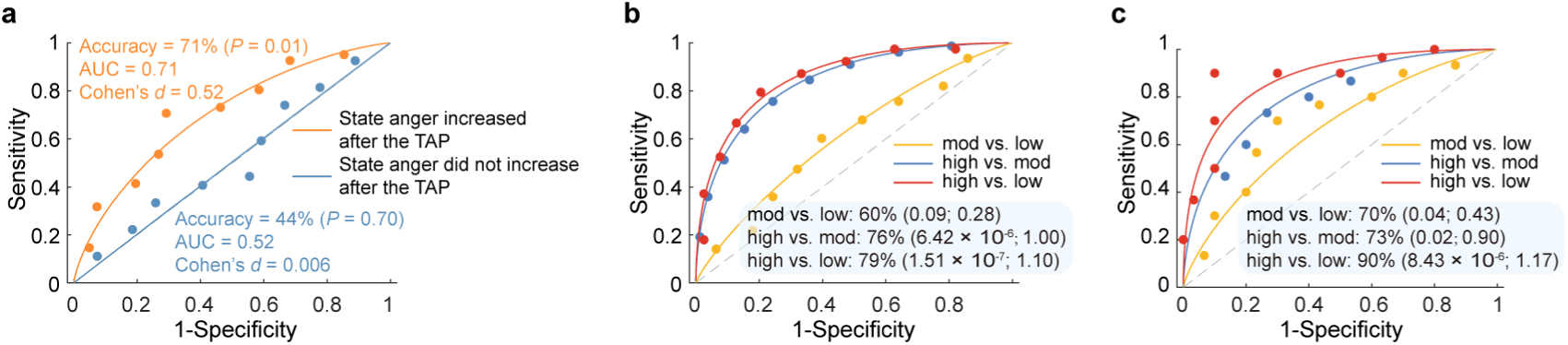
Neural evidence for anger-and disgust-related components of outrage. **a**, The MONS accurately discriminated aggression following high versus low provocation among participants who exhibited increased post-task state anger, but not among those without increased post-task state anger. **b,c**, The MONS accurately distinguished different levels of subjective disgust in both the disgust discovery (**b**) and validation (**c**) cohorts, except for the moderate-versus-low comparison in the discovery cohort (shown as forced-choice classification accuracy, *P* value and Cohen’s *d*). *P* values were based on two-sided independent binomial tests.

### Separable signatures of outrage, general negative affect and affective arousal

As previous theory-based and behavioral studies have suggested conceptual commonalities as well as distinctions between outrage and broader affective constructs such as negative affect and affective arousal^3,15,17^, we next examined to what extent these emotional experiences share or exhibit dissociable neurofunctional representations. We compared the MONS with the established neural signatures of negative affect (MNAS^47^) and affective arousal (BAAS^48^) across spatial and functional dimensions.

On the spatial level, the three signatures showed partially overlapping but distinct regional contributions. The MONS showed preferential contributions from the PCC, whereas several regions, including the ACC, MCC, precuneus and anterior insula, contributed to both the MONS and MNAS. Other regions, including the hypothalamus, PAG, thalamus, dMPFC, middle insula, TPJ, vLPFC, dLPFC and ventral striatum, showed reliable contributions across all three signatures, although their relative contributions differed between signatures (Fig. 6a,b; Supplementary Results and Supplementary Table 7).

**Fig. 6.**
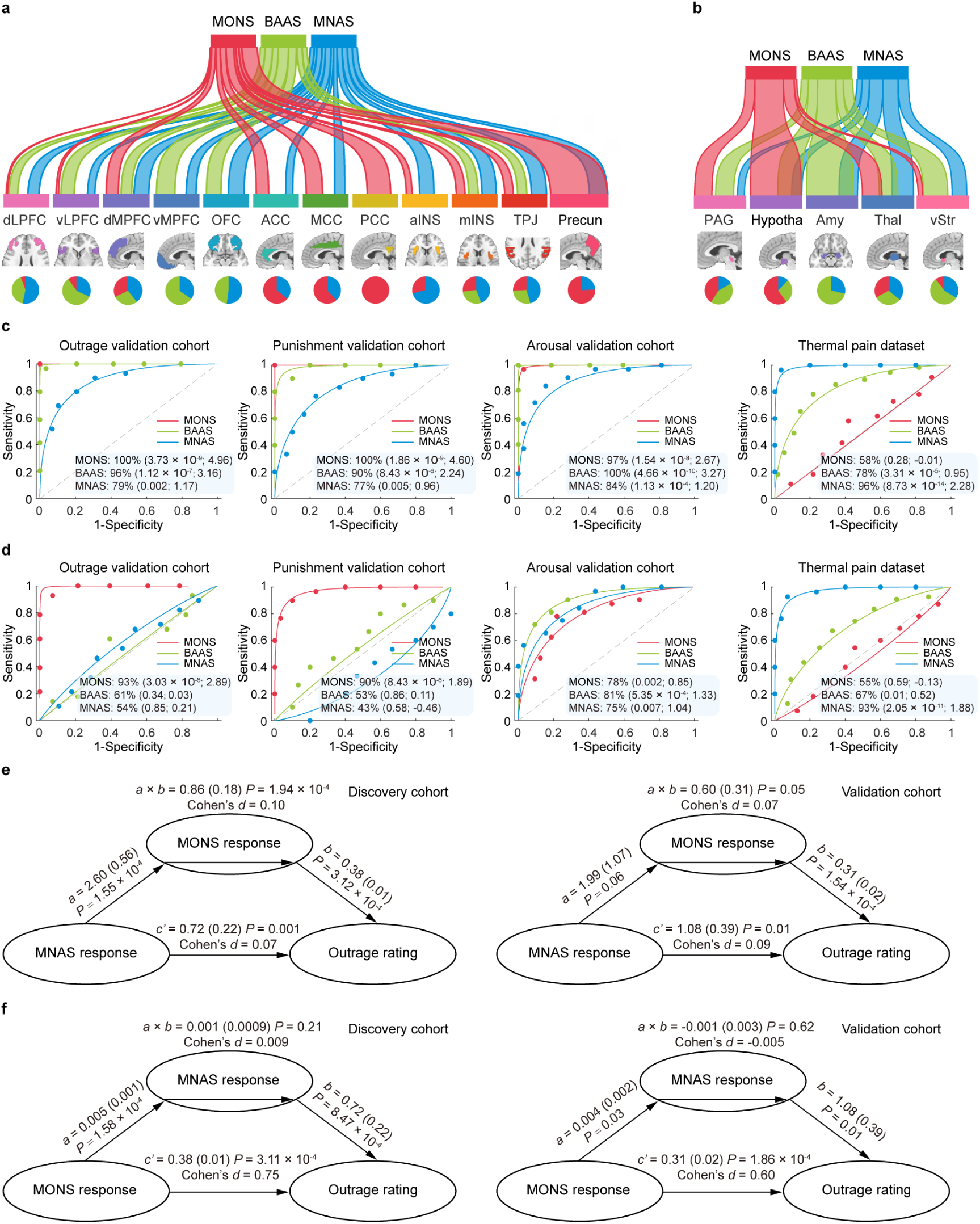
Comparing neurofunctional signatures for outrage, general negative affect and affective arousal (MONS, MNAS, BAAS). **a,b**, River plots showing spatial similarity (cosine similarity) between stable decoding maps and a priori ROIs. Ribbons are normalized by the maximum cosine similarity across all cortical ROIs (**a**) and subcortical ROIs (**b**). Stable decoding models were thresholded at FDR *q* < 0.05, and similarity analyses were restricted to positive voxels only. Ribbon locations in relation to the boxes are arbitrary. Pie charts show the relative contributions of each model to each ROI (i.e., the percentage of voxels showing the highest cosine similarity with each map). **c,d**, MONS, MNAS and BAAS more accurately classify their respective targeted domains during high versus low (**c**) and high versus moderate (**d**) intensity discrimination (shown as forced-choice classification accuracy, *P* value and Cohen’s *d*). *P* values were derived from two-sided binomial tests. **e**, MONS response partially mediates the association between MNAS response and subjective outrage ratings in both the discovery and validation cohorts. **f**, MNAS response does not significantly mediate the effect of MONS response on outrage ratings in either the discovery or the validation cohort. **e,f**, Two-sided *P* values were derived from bootstrap tests with 10,000 samples. vMPFC=ventromedial prefrontal cortex; vStr=ventral striatum.

On the functional level, the MONS showed greater sensitivity to outrage and punishment than the MNAS and BAAS, with substantially larger effect sizes for high-versus-low intensity discrimination (1.57–4.24-fold for outrage and 2.05–4.79-fold for punishment; Fig. 6c). Conversely, the MNAS and BAAS showed greater sensitivity to their respective target emotional domains. Additional high-versus-moderate intensity analyses further supported the relative specificity of the MONS for outrage and punishment, as indicated by 13.76–96.33-fold and 4.11–17.18-fold larger effect sizes for outrage and punishment, respectively, than those of the MNAS and BAAS (Fig. 6d). Moreover, consistent with the findings that the MONS did not respond to thermal pain at either high-versus-low or high-versus-moderate intensity discrimination (Fig. 6c,d), it failed to significantly predict responses in additional negative-affective contexts^47^, including aversive images, mechanical pain and aversive sounds (Supplementary Fig. 7).

Finally, multilevel mediation analyses provided further evidence for partially overlapping but dissociable representations. The MONS response significantly mediated the relationship between MNAS/BAAS responses and outrage ratings, whereas the reverse mediation was weaker or non-significant (Supplementary Results, Fig. 6e,f and Supplementary Fig. 8). Similar results were observed when punishment ratings were examined as the outcome (Supplementary Results and Supplementary Fig. 9,10).

Importantly, the MONS also retained its predictive utility for subjective outrage after controlling for BAAS-derived arousal-related variance (Supplementary Methods, Supplementary Results and Supplementary Fig. 11).

Together, these findings indicate that subjective outrage shares partially overlapping neurofunctional representations with general negative affect and affective arousal, while retaining greater sensitivity to outrage-related experience and punishment processing.

### Validating the outrage signature in traditional moral judgment contexts

Previous studies have shown that outrage is strongly associated with moral judgment^35,50,52,53^, and our pre-rating experiment supported this relationship, revealing a stimulus-level correlation of *r*=0.66±0.24 between outrage and perceived moral wrongness (Supplementary Table 2). We thus examined whether the neural signature of subjective outrage could predict perceived moral wrongness across six independent moral judgment datasets that applied traditional moral judgment paradigms^54^, including brief vignettes (*“You see a man leaving his family business to go work for their main competitor”*) (four datasets, studies 8–11; *n*=64, 30, 27 and 30, respectively; Supplementary Table 5) and pictures depicting immoral situations (two datasets, studies 12–13; *n*=30 and 30, respectively; Supplementary Table 5). To assess the predictive performance of the MONS while reducing the potential contribution of sensory processing associated with the audiovisual training stimuli, we evaluated the signature excluding the visual and auditory cortices. Across the four vignette-based datasets, the MONS accurately predicted moderate versus low moral wrongness (accuracies=[65%, 74%], *P*<0.05, Cohen’s d=[0.52, 0.76]), but did not significantly discriminate moral wrongness in either the high-versus-low or high-versus-moderate contrasts (Fig. 7a). Consistent with this limited predictive performance, overall and within-subject correlations between signature responses and moral wrongness ratings were weak and nonsignificant across the four vignette-based datasets (Supplementary Table 8). In the two picture-based datasets, the MONS did not significantly predict either the immoral-versus-neutral or the immoral-versus-moral contrast (Fig. 7b). Results obtained using the whole-brain MONS were similarly inconsistent across the six datasets (Supplementary Results, Supplementary Fig. 12a,b and Supplementary Table 9). Notably, a few contrasts reached significance in the opposite direction (Supplementary Fig. 12), which may reflect contributions from visual and auditory cortices. Collectively, these results suggest that the neural representation of outrage induced by short social-media videos is distinguishable from moral judgment as assessed by traditional experimental paradigms.

**Fig. 7.**
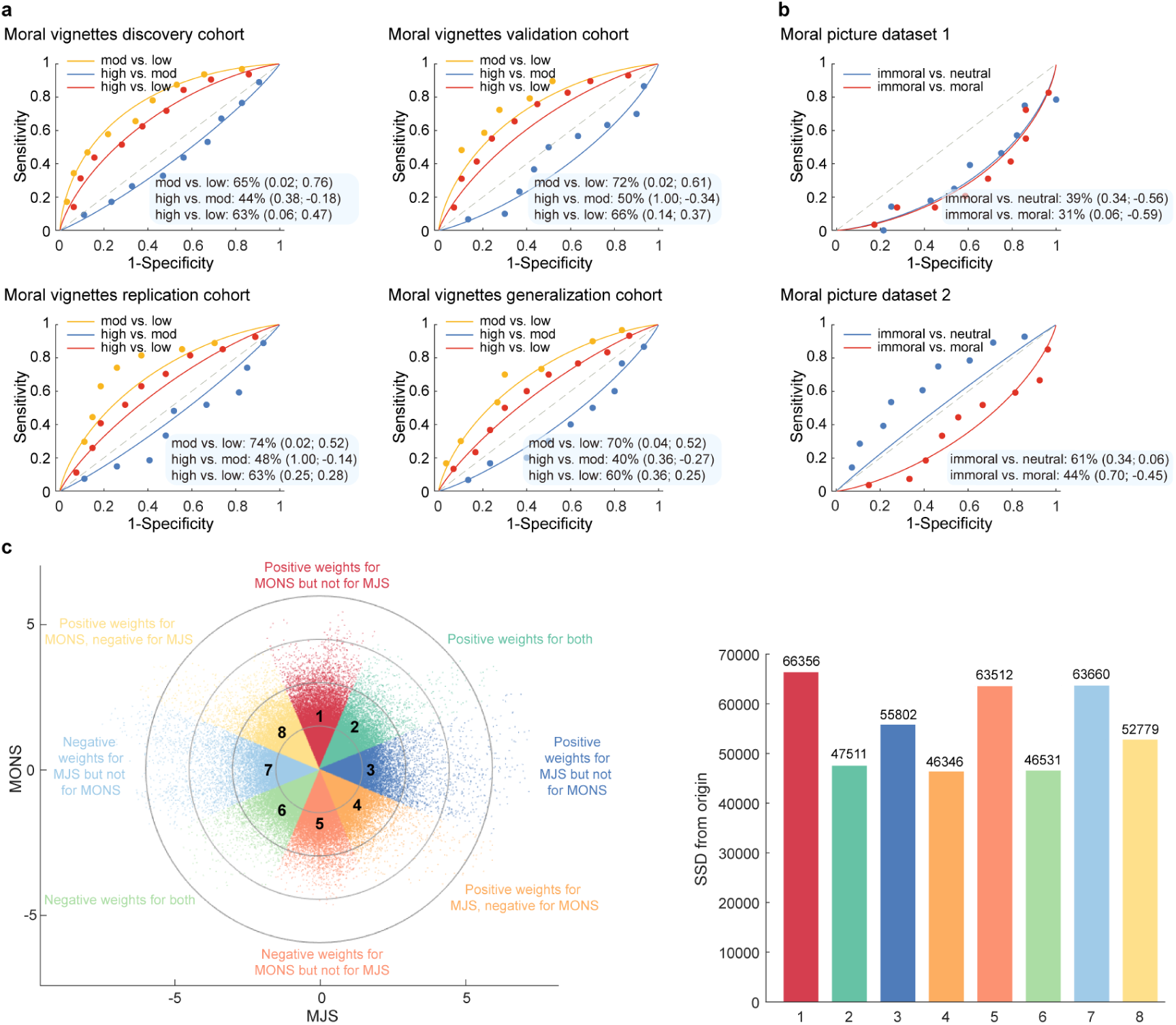
Validation of the MONS excluding the visual and auditory cortices in traditional moral judgment contexts and voxel-level spatial similarity between MONS and MJS. **a**, The MONS excluding the visual and auditory cortices was not sensitive to distinguish different levels of moral wrongness across the four vignette-based moral judgment datasets, except for the comparison between moderate and low moral wrongness contrast (*P*<0.05) (shown as forced-choice classification accuracy, *P* value and Cohen’s *d*). **b**, The MONS inaccurately discriminated between immoral and moral or neutral conditions in two picture-based moral judgment datasets. **a,b**, *P* values were based on binomial tests, two-sided. **c**, Scatter plot displaying normalized voxel weights for MONS (*y*-axis) and MJS (*x*-axis). Each dot represents a voxel. Bars on the right show the sum of squared distances from the origin (0, 0) in each octant. Different colors are assigned to the eight octants that reflect voxels of shared positive or shared negative weights (Octants 2 and 6, respectively), uniquely positive weights for the MONS (Octant 1) or for MJS (Octant 3), uniquely negative weights for the MONS (Octant 5) or MJS (Octant 7), and voxels with opposite weights for the two neural signatures in the remaining octants (Octants 4 and 8).

To further determine the spatial relationship between the neural representations of subjective outrage and moral judgment in static or hypothetical contexts, we assessed the whole-brain spatial correlation between MONS and the moral judgment signature (MJS; developed in ref.^54^; Supplementary Methods). The pattern weights of the two signatures were not significantly correlated on the whole-brain level (*r*=-0.02, permutation test one-tailed *P*=0.23). We next examined their joint distribution to characterize the spatial correspondence between their normalized voxel weights, plotting MONS on the *y*-axis and the MJS on the *x*-axis (Supplementary Methods; for a similar approach see ref.^77^). As shown in Fig. 7c, stronger weights across the whole brain (as indicated by higher sum of squared distances (SSD) from the origin) were observed in the nonshared Octants (1, 3, 5, 7).

Overall, these findings suggest separable neural representations of naturalistic social-media-induced outrage and the mental processes induced by traditional moral judgment paradigms, which may reflect the explicit cognitive evaluation of moral violations and lack of a strong emotional component in the latter.

## Discussion

As a powerful driver of attention, engagement and information diffusion, outrage has become increasingly influential in shaping online interactions and behavioral responses^1,3,6^. Our current understanding of outrage in the social media age is largely based on theoretical accounts (e.g., refs.^1,4,6,14,17^) and large-scale behavioral studies (e.g., refs.^2,3,5,23,25,31^), providing important insights into how social media shapes emotions and behavior. However, how the brain responds to an informational ecosystem saturated with “***rage bait***”^36^ remains largely unexplored. Here, using naturalistic fMRI during massive exposure to diverse short-form videos depicting moral violations from social media, combined with predictive modeling, we identified and validated a sensitive and robust neural signature of subjective outrage (MONS). The MONS accurately predicted momentary outrage experience and punishment tendencies on both population and individual levels. Subjective outrage was represented in distributed cortical and subcortical systems, with no isolated region or network achieving the predictive performance of the whole-brain model. The MONS further predicted subjective anger and disgust, providing neurobiological evidence for the prevailing model that outrage reflects a blend of anger and disgust^2–5,13,50,51^. Multiple analyses demonstrated that outrage was not reducible to broader affective processes. Relative to established neural signatures of general negative affect (MNAS^47^) and affective arousal (BAAS^48^), the MONS exhibited dissociable spatial representations, greater sensitivity to outrage (and punishment tendencies) and retained predictive utility after controlling for affective arousal. Moreover, the MONS robustly mediated the response of the other two models (MNAS, BAAS) on both subjective outrage and punishment ratings, but not vice versa. Notably, however, the MONS did not reliably track perceived moral wrongness in static and hypothetical moral judgment contexts which emphasized explicit cognitive evaluation of moral violations. Together, we identified a robust, distributed and functionally specific neural signature of moral outrage induced by naturalistic social media.

The fact that outrage can motivate punishment of transgressors has been well documented in previous studies^6,7,10,50^, with such punishment potentially contributing to the enforcement of social norms and the maintenance of pro-sociality^10–12,78,79^. Responding to calls to examine outrage intensity^80^ and social punishment in more dynamic settings beyond static paradigms^81^, we simultaneously assessed ratings of outrage experience and punishment tendencies while participants viewed ecologically valid short-form videos showing moral transgressions in the scanner. We found that trial-by-trial fluctuations in punishment ratings closely tracked outrage ratings, such that greater experienced outrage was associated with harsher punishment decisions. Converging neural evidence was observed, as the MONS accurately predicted punishment tendencies, providing neurobiological support for the link between experienced outrage and ensuing punitive responses. The close correspondence between outrage and punishment may partly reflect their immediate succession in our paradigm, which resembles the rapid-response dynamics of social media, where users can like, react to or disseminate content almost instantaneously^4,6^. This may explain the strong behavioral motivation and attentional capture induced by outrage content on social media, facilitating online consumption and sharing^18,25^. Combined, the behavioral and neural findings indicate that outrage is closely linked to subsequent punishment tendencies, particularly in contexts that favor swift responses.

Consistent with growing evidence that emotions are represented in distributed brain systems^45–49^, our findings reveal that subjective outrage emerges from coordinated activity across subcortical and cortical systems implicated in threat detection (PAG, brainstem, hypothalamus and thalamus)^9,61,62^, social cognition (e.g., mentalizing/theory of mind; precuneus, PCC, MCC, dMPFC, TPJ, superior temporal sulcus, superior temporal gyrus, middle temporal gyrus, temporal pole)^9,62–67^, conflict detection and self-regulatory control (ACC, inferior frontal gyrus)^62,68,69^, interoception (posterior insula)^70,71^, approach-related processing (caudate)^62^, social aversion to norm violations (fusiform gyrus)^81^ and motor preparation and action implementation (supplementary motor area, precentral gyrus, inferior parietal lobule, supramarginal gyrus, substantia nigra and cerebellar regions)^9,72–74^. Importantly, an independent functional decoding analysis based on the BrainMap database converged with this interpretation by linking representative MONS subregions primarily to social cognition, negative affect, arousal/salience, domain-general cognition and action-related domains, which dovetails with constructionist theories of emotion proposing that emotional experiences arise from interactions among multiple systems supporting affective, sensory, cognitive and motor functions^82^. Moreover, comparisons with models based on localized regions and large-scale networks showed substantially lower predictive performance than the whole-brain MONS, further supporting a distributed rather than localized neural representation of subjective outrage (Supplementary Results). This distributed organizational principle accords with appraisal^83^ and constructionist^84^ theories of emotion, which propose that subjective emotional experiences are implemented by distributed neural systems rather than localized brain modules.

Although outrage is negatively valenced and highly arousing^17^, our findings demonstrate that its neural representation is dissociable from established predictive models of general negative affect (MNAS^47^) and affective arousal (BAAS^48^). Spatial similarity analyses revealed differential contributions of brain systems supporting emotional awareness and appraisal, social cognition, cognitive control and threat detection across the three signatures. Regions implicated in mentalizing (e.g., precuneus, PCC)^62^, conflict detection and self-regulation (e.g., ACC)^62,68,69^, and threat detection (e.g., hypothalamus)^61^ contributed more strongly to the MONS than to the MNAS or BAAS. By contrast, lateral and ventromedial prefrontal regions (dLPFC, vLPFC and vMPFC) contributed comparably little to the MONS. Given the roles of these regions in behavioral control and emotion regulation^69,85^, these findings may suggest that subjective outrage relies relatively less on top-down regulatory processes. The absence of stable positive contributions from the OFC and amygdala, which support valuation and behavioral flexibility^86,87^, should be interpreted cautiously given their negative MONS weights (Supplementary Results). The ventral striatum contributed less strongly to the MONS than to the MNAS and BAAS. Given its well-established role in reward processing and incentive motivation^88,89^, this relative difference is notable in light of the remarkable capacity of outrage-eliciting content to attract sustained engagement on social media^1,4,8^. Recent behavioral evidence similarly suggests that although users frequently engage with outrage-related content, they simultaneously report preferring lower levels of outrage in their online feeds^90^.

The following functional similarity analyses showed that the MONS was more sensitive than the MNAS in predicting both subjective outrage and punishment tendencies, while not reliably predicting general negative affect. Mediation results corroborated these findings, showing that the MONS response mediated the association between MNAS response and both subjective outrage and punishment ratings, but not vice versa. These findings conceptually replicate previous work demonstrating that a social-media-text-based outrage classifier exhibited discriminant validity relative to a text-based classifier trained to capture broader negative affect^3^. The BAAS predicted both outrage and punishment at the low-to-high intensity range, but not at the moderate-to-high intensity range. Parallel to this observation, mediation analyses revealed reciprocal mediation between BAAS and MONS responses; however, the mediation effect through the MONS was substantially stronger than that through the BAAS. Furthermore, the MONS retained robust predictive performance for subjective outrage after controlling for affective arousal. These findings converge with behavioral evidence showing that the viral spread of moral-emotional content, including outrage, cannot be fully explained by the extent to which such content is arousing^15^. Altogether, these findings suggest that subjective outrage shares affective components with both general negative affect and affective arousal while remaining neurofunctionally distinguishable from both, potentially providing a neural basis for its distinctive impact on attention, engagement and moral behavior in online environments.

Of note, the neural signature of outrage failed to reliably capture perceived moral wrongness across all six independent moral judgment datasets^54^, despite consistent evidence from previous conceptual and behavioral studies^35,50,52,53^—as well as our own pre-rating experiment—that the two are closely correlated. This limited generalization, together with the distinct spatial distributions of MONS and the moral judgment signature, suggests that naturalistic outrage and moral judgment elicited by traditional paradigms in hypothetical or static contexts may involve separable neural representations. The classical moral judgment paradigms were based on written vignettes or static pictures^54^ (see also ref.^37^), which primarily emphasize explicit cognitive evaluation of moral violations. In contrast, our short-form videos were designed to elicit affectively engaging moral outrage in naturalistic social-media contexts. Such naturalistic contexts may offer greater ecological validity for investigating morality in the digital age, where outrage-eliciting content appears to be particularly prevalent and potent^6^. Online platforms are also increasingly shaping the rapid cultural evolution of social norms and normative thinking^21^. The growing prominence of social media may therefore make emotionally engaging, context-rich moral experiences an increasingly important complement to the explicit evaluation of moral violations typically examined in classical paradigms. To our knowledge, the present study is among the first to leverage naturalistic social-media short-form videos to examine how outrage is represented in the brain. These findings highlight the need to extend established approaches to moral neuroscience by incorporating naturalistic paradigms that capture affectively engaging moral experiences, thereby providing a more comprehensive account of the neural processes underlying outrage in contemporary social contexts.

The present study has several limitations. First, whereas previous studies have focused on the expression of emotions—for example, sharing text/commentary containing moralized and emotional language (e.g., outrage; see refs.^3,5,15,25,35,51^)—we examined the subjective experience of outrage. This focus represents a strength of our outrage-induction fMRI paradigm because subjective emotional experience is difficult to infer directly from online behavioral data^18^. Although expressing outrage online is one possible behavioral response associated with experiencing outrage^80^ and the two are expected to be highly correlated^3^, recent causal evidence further supports this relationship by showing that moral transgressions induce outrage, which subsequently promotes preferences for sharing outrage-eliciting content^31^. Whether the MONS generalizes to online expressions of outrage, such as sharing, remains unknown. Future studies could examine how the neural representation of outrage experience relates to its online expression and the spread of moral information, particularly given evidence that misinformation exploits outrage to spread online^2,14^. Second, although our paradigm captured rapid exposure and responding to morally violating content^4,6^, it did not reproduce the interactive features of social media, such as scrolling, liking, sharing, and commenting, or publicly sanctioning transgressors. Thus, punishment ratings provided an experimentally tractable index of punishment tendencies rather than a direct measure of online punitive behavior. Because online punishment is often public, highly visible and low-cost^1,6,8^, the present data cannot determine whether the punishment tendencies predicted by the MONS would translate into online behaviors such as public shaming, pile-ons, or conflict escalation^6,8,24^. Future work may combine neural measures with more interactive social-media environments to determine how outrage translates into punishment-related behaviors. Thirdly, our experimental stimuli were drawn from short-form video platforms (e.g., Douyin) and presented within an East Asian cultural context, potentially limiting generalizability across platforms and cultures. Future work may test the MONS across more diverse platforms and cultural contexts.

In conclusion, the present study developed a whole-brain neural signature of experienced outrage. The MONS sensitively predicted momentary outrage experience and punishment tendencies. Subjective outrage was encoded in distributed brain systems rather than isolated regions. The MONS also captured subjective anger and disgust. However, the MONS was distinct from the neural representations of general negative affect and affective arousal, and did not predict moral wrongness in static and hypothetical moral judgment contexts, indicating a more affective state of moral processing in outrage. Taken together, these findings represent an important step toward understanding the neurobiological basis of subjective outrage in context with social media.

## Methods

### Short-form video clips and pre-rating procedure

The short-form video clips were drawn from a large pool of materials downloaded from Douyin (the Chinese equivalent of TikTok), WeChat (a social communication application) and Kuaishou (another popular short-form video application in China). This selection process was conducted by two researchers (the first and fifth authors). An initial set of 120 audiovisual clips (each approximately 25 s long) was obtained, all depicting real-world situations. The clips were divided into two subsets, and two independent groups of participants (*n*=21 per group, with 11 females in each group; group 1: mean± s.d. age=20.43±2.15 years; group 2: mean± s.d. age=20.29±2.31 years) were recruited to attentively watch one subset.

Following a validated pre-rating procedure^48^, after watching each audiovisual clip, participants rated their levels of outrage, outrage consistency, arousal, perceived moral wrongness, and intended punishment toward the transgressor(s) (if any) depicted in the clip using 9-point Likert scales (1=neutral/slightest outrage or lowest consistency or lowest arousal or not morally wrong or no punishment; 9=strongest outrage or highest consistency or highest arousal or extremely morally wrong or severest punishment). Of note, outrage consistency was operationalized as the intensity of the elicited outrage, where level 1 indicated that the intensity of the elicited outrage was very inconsistent throughout the clip and 9 indicated that the intensity of the elicited outrage was very consistent throughout the clip. Before the pre-rating experiment, participants were presented with illustrative examples of punishment adapted from previous studies^91,92^ (e.g., “I would hit, shove, insult or yell at the transgressor(s) in the clip”) to promote a common understanding of the punishment construct. The pre-rating experiment was implemented in PsychoPy^93^ (v3.2.4) on a Windows platform. Based on the rating data, 45 highly outrage-inducing clips (outrage consistency>7; Supplementary Table 1) were selected as outrage-inducing stimuli for the fMRI experiment. These stimuli include a wide range of diverse moral transgressions, such as uncivil behavior, cat/child/elderly abuse, domestic violence, obstruction of public duty, intergroup conflict, genocide, etc.

In addition, another 15 low-arousing neutral clips were selected from previous studies^48,94^. These stimuli depict everyday social scenarios, such as getting a haircut or shave at a barbershop, spending time with a pet dog, receiving a massage at a massage parlor, and elderly people practicing Tai Chi, etc.

The 45 outrage-inducing clips from the pre-rating procedure, together with 15 neutral clips, constituted the final stimulus set used in study 1. All 60 clips were embedded in an East Asian cultural context.

### Participants in the discovery cohort (study 1)

A total of 83 participants were recruited from the University of Electronic Science and Technology of China (UESTC) for the discovery cohort (study 1). All participants were right-handed, had normal or corrected-to-normal vision, reported no history of mental or physical disorders, no MRI contraindications, and were free of current or habitual substance or medication use. Following data quality control, three participants (one female) were excluded due to excessive head motion during fMRI scanning (>3 mm), yielding a final sample of *n*=80 participants (40 females; mean±s.d. age=21.89±2.45 years).

### Stimuli and paradigm used in the discovery cohort (study 1)

The experiment comprised 4 runs, each including 15 clips. Participants were instructed to pay attention to the audiovisual stimuli and naturally experience the elicited emotion. Each trial consisted of a ∼25 s clip, followed by a jittered fixation cross (1-1.2 s). Participants then had 4 s to report their experienced level of outrage on a 9-point Likert scale (1=neutral/slightest outrage; 9=strongest outrage). After a jittered fixation cross epoch (1-1.2 s), participants rated their willingness to punish the transgressor(s) (if any) in the clip using another 9-point Likert scale (1=no punishment; 9=severest punishment), also within 4 s. Each trial ended with a jittered fixation cross epoch (10-12 s) serving as a washout period before the next clip (Fig. 1a). Stimulus presentation and behavioral data collection were controlled using MATLAB 2015a (MathWorks, Natick, MA) and Psychtoolbox (http://psychtoolbox.org/). Before the formal fMRI session, participants completed a practice session outside the scanner to familiarize themselves with the procedure. The same illustrative punishment examples^91,92^ used in the pre-rating procedure were presented to ensure a consistent understanding of the construct.

Notably, the outrage-eliciting video clips selected through the pre-rating procedure evoked predominantly high levels of outrage and punishment during the fMRI experiment. Specifically, participants’ ratings were largely concentrated toward the upper end of the scale (i.e., scores of 6– 9), whereas lower ratings (i.e., 2–5) were comparatively infrequent. This distribution provides indirect support for the effectiveness of the pre-rating procedure, as reflected in the high mean ratings of outrage (mean ± s.d.=7.35±0.61) and punishment (mean ± s.d.=7.44±0.88) across the 45 selected outrage-inducing clips (Supplementary Table 1). To ensure a sufficient number of trials within each condition for subsequent analyses, we applied a binning procedure to the rating scale. Ratings of 1–3 were combined into level 1, ratings of 4–5 into level 2, and ratings of 6–7 into level 3, while ratings of 8 and 9 were treated as levels 4 and 5, respectively (see Results for more details).

### MRI data acquisition and preprocessing in the discovery cohort

MRI data were collected using a 3T scanner (GE MR750; General Electric Medical System, Milwaukee, WI, USA) (see Supplementary Methods for details). Functional MRI data were preprocessed and analyzed using SPM12 (Statistical Parametric Mapping; https://www.fil.ion.ucl.ac.uk/spm/software/spm12/), following procedures consistent with our previous studies^45,46,95^. The first five volumes of each functional run were discarded to allow for signal equilibration. Before preprocessing, motion-related outliers were identified for each run based on either of the following criteria: (1) signal intensity larger than three s.d. from the global mean or (2) signal intensity and Mahalanobis distances larger than ten mean absolute deviations (https://github.com/canlab/CanlabCore). Each timepoint identified as outliers was subsequently incorporated into the first-level design matrix as a separate nuisance regressor.

Preprocessing included slice-timing correction and realignment to correct for head motion, followed by unwarping to account for magnetic field inhomogeneities. High-resolution anatomical images were segmented into gray matter, white matter, cerebrospinal fluid, bone, fat and air. The resulting skull-stripped and bias-corrected structural images were co-registered to the functional images. Functional images were then normalized to the Montreal Neurological Institute (MNI) template space and resampled to 2×2×2 mm³ isotropic voxels, and subsequently smoothed using an 8-mm full-width at half maximum (FWHM) Gaussian kernel.

### First-level fMRI analysis in the discovery cohort

Given that the primary goal of the present study was to develop a neural signature of subjective moral outrage experience (i.e., MONS), and that outrage and punishment ratings were highly correlated and exhibited closely aligned trial-wise profiles (see Results), first-level analyses were designed to model outrage-related neural responses. For each participant, two separate univariate general linear models (GLMs) were specified at the first level. The first GLM was designed to generate beta maps for subsequent prediction analyses. In this model, neural responses to the clip presentation period were modeled using five condition-specific regressors corresponding to the binned outrage ratings (levels 1–5), allowing brain activity associated with each rating level to be estimated separately. In addition, two boxcar regressors (4 s) were included to capture activity during the two rating periods (i.e., outrage and punishment rating phases, respectively), thereby accounting for motor-related processes. Fixation periods were treated as the implicit baseline. The second GLM adopted a parametric modulation approach to examine rating-dependent neural responses. This model included a regressor corresponding to the clip presentation period, as well as separate regressors modeling the two rating phases. Trial-wise outrage ratings (levels 1–5) for each clip were entered as a parametric modulator of the clip presentation regressor, enabling identification of brain regions in which activity scaled with subjective outrage intensity.

All task-related regressors were convolved with the canonical hemodynamic response function. Functional runs were concatenated within each participant using the SPM function spm_fmri_concatenate.m. Nuisance regressors included: (1) run-specific intercepts to account for mean signal differences across runs; (2) head motion parameters (three translations and three rotations), along with their squares, derivatives and squared derivatives (24 regressors in total); and (3) regressors indexing time points identified as motion-related outliers.

### Participants in the validation cohort (study 2)

To evaluate the performance of the outrage signature (MONS) developed using the discovery cohort (i.e., study 1), an independent sample was collected as a validation cohort (study 2) using the same experimental paradigm. A total of 32 healthy participants were recruited from UESTC. Enrollment criteria were identical to those applied in study 1. After quality control, two male participants were excluded due to excessive head motion during fMRI acquisition (>3 mm), resulting in a final sample of *n*=30 participants (16 females; mean±s.d. age=20.80±2.67 years).

### Stimuli, paradigm, MRI data acquisition, preprocessing and first-level fMRI analysis in the validation cohort (study 2)

The validation cohort (study 2) followed the same experimental procedures and data processing pipelines as the discovery cohort (study 1), including stimuli, task paradigm, rating binning, MRI data acquisition, preprocessing and participant-level GLM specification. To further characterize the relationship between outrage and punishment from the perspective of predictive modeling, an additional GLM was constructed to derive beta estimates associated with binned punishment ratings (levels 1–5) (see Supplementary Methods).

### MVPA

We used whole-brain multivariate pattern analysis^45–48,58^ to identify a distributed pattern of weighted brain activity that best predicted the self-reported outrage ratings. Specifically, we employed the SVR algorithm (linear kernel with *C*=1) implemented in the Spider toolbox (http://people.kyb.tuebingen.mpg.de/spider), where beta images corresponding to each rating level served as input features and the self-reported outrage ratings were used as outcome variables. Whole-brain data masked with a gray matter mask from the discovery cohort were used. To test the performance of the outrage-predictive pattern and to rule out the possibility of model overfitting^58,96,97^, we implemented a rigorous 10 repeats of 10-fold (i.e., 10×10-fold) cross-validation procedure where participants in the discovery cohort were randomly partitioned into 10 folds (8 participants per fold) using MATLAB’s cvpartition function. For each fold, the model was trained on data from 72 participants and tested on the remaining 8 participants. This procedure was iterated until each fold had served as the test set once, and the entire process was repeated 10 times with different random partitions to obtain a stable estimate of prediction performance and minimize potential biases associated with data splitting^45,46,95^.

To evaluate the predictive performance of the neurofunctional signature of outrage, we assessed both the correspondence between predicted and observed outrage ratings and the magnitude of prediction error. Pearson correlations (*r*) between the cross-validated predictions and the true ratings were calculated at both the overall level (between- and within-subjects; 370 pairs in total) and within-subject level, serving as indices of effect sizes. These correlations quantified the correspondence between predicted and observed variation in subjective outrage. In addition, RMSE was computed to index the magnitude of prediction error. Thus, *r* and RMSE provided complementary measures of prediction performance, with *r* capturing correspondence in variation and RMSE quantifying absolute prediction error (for similar model evaluation metrics, see ref.^47^). To further characterize the discriminative performance of the outrage signature trained on the discovery cohort, we conducted forced-choice classification analyses based on receiver operating characteristic (ROC) curves. Classification accuracy was assessed for each adjacent pair of outrage rating levels (2 versus 1, 3 versus 2, 4 versus 3, and 5 versus 4) as well as for broader pairs (high (ratings 4–5) versus low (rating 1), high versus moderate (ratings 2–3), and moderate versus low). In each pair, model (pattern) response values for two conditions within the same participant were compared and the higher response was selected as more outraged, yielding a ‘threshold-free’ accuracy measure. Statistical significance of the classification accuracies was evaluated using two-sided binomial tests against the chance level (50%).

To further assess how well the MONS predicted outrage (as well as punishment tendency) on datasets other than the discovery cohort and rule out overfitting^96^, the MONS was applied to independent datasets. These included the validation cohort (study 2; the outrage condition) and the validation cohort (study 2; the punishment condition). For each dataset, pattern responses were computed as the dot product between the MONS weight map and each GLM-derived fMRI activation map.

### Within-subject trial-wise prediction

To determine whether the MONS could predict moment-to-moment fluctuations in subjective outrage on the single-trial level, we conducted a trial-wise analysis based on single-trial fMRI estimates^45,46,98,99^. Specifically, a separate GLM was constructed for each participant in which each clip presentation period was modeled with an individual regressor, thereby yielding trial-specific beta estimates. All task-related regressors were convolved with the canonical hemodynamic response function, and nuisance regressors were identical to those included in the primary first- level analyses. Single-trial beta estimation can be particularly susceptible to noise contamination arising from transient artifacts, such as head motion or scanner pulse artifacts occurring during individual trials. To improve the reliability of trial-wise estimates, variance inflation factors (VIFs) were computed for each trial-specific regressor as an index of design-related collinearity with nuisance regressors and neighboring task regressors. Trials with VIF values exceeding 3 were excluded from subsequent analyses. Using this criterion, approximately 8.9% and 8.6% of trials were excluded from the discovery and validation cohorts, respectively. The resulting single-trial beta maps were then submitted to MONS pattern expression analysis by calculating the dot product between the vectorized activation maps and the MONS weight map. For subjects in the discovery cohort, the same 10×10-fold cross-validation procedure described above was used to obtain the MONS response of each single-trial beta map for each subject. We further evaluated trial-wise prediction performance in the validation cohort by applying the MONS trained on the discovery cohort to single-trial activation maps derived from the validation cohort. Prediction performance was assessed for both outrage ratings and punishment ratings to examine whether the MONS extended beyond subjective outrage to capture closely related punishment responses on the trial-by-trial level.

### Determining brain regions associated with and predictive of subjective outrage

To characterize the neural substrates underlying the subjective experience of outrage, we employed a series of analyses on the population-level outrage signature (MONS). First, bootstrap tests were performed to identify brain regions that contributed stably to prediction performance across participants. Specifically, 10,000 bootstrap samples were generated from the discovery cohort using sampling with replacement, and the prediction model was re-estimated for each sample. Voxel-wise *z*-scores and corresponding two-tailed uncorrected *P* values were then derived from the mean and s.d. of the resulting bootstrap distributions. Brain regions exhibiting stable predictive weights after correction for multiple comparisons (FDR *q*<0.05) were considered to show reliable contributions to the MONS^60^. Second, following previous neurofunctional signature studies^45–47^, we transformed the population-level MONS pattern (backward model) into ‘activation pattern’ (forward model), which maps individual voxels to the response (fitted values) in the multivariate model, reflecting the association between voxel-wise activity and the model-predicted response^55^. In the statistical literature, this reconstructed ‘activation pattern’ is also referred to as ‘structure coefficient’^100,101^. Reliable outrage predictive and associative regions of the MONS were identified using the one-sample *t* test, thresholded at FDR *q*<0.05 corrected for multiple comparisons. Core regions underlying subjective outrage were defined as having voxels that were consistently identified in both the backward predictive-weight map and the forward activation pattern map, thereby reflecting regions that were both strongly predictive of and reliably associated with subjective outrage experience. For methodological details regarding the identification of brain regions associated with subjective outrage on the individual level, see Supplementary Methods.

Next, we examined whether the subjective experience of outrage could be sufficiently predicted from localized neural systems rather than the whole-brain model. To address this question, we conducted whole-brain searchlight-based (three-voxel-radius spheres) and atlas-based parcellation (279-region version of the Brainnetome Atlas that additionally incorporated brainstem, midbrain, and cerebellum regions^75,76^) analyses to identify local regions predictive of outrage, and compared their prediction performance with that of the whole-brain MONS model. In addition, we compared the prediction performance of a set of ROIs previously documented in outrage-related processing, social cognition, and threat detection^9,61,62,64–66^ to the whole-brain model. Cortical ROIs included the OFC, ACC, MCC, PCC, anterior insula, middle insula, TPJ, and precuneus, whereas subcortical ROIs included the PAG, hypothalamus, amygdala, and thalamus. We further compared the prediction performance of large-scale networks to that of the MONS. The networks of interest comprised seven large-scale cerebral networks^102^ and a subcortical network (including the striatum, thalamus, hippocampus, and amygdala). For each searchlight sphere, atlas-defined parcellation, ROI, or large-scale network, separate prediction models were trained and tested using the discovery cohort (10×10-fold cross-validation), and their predictive performance was subsequently evaluated in the independent validation cohort.

Additionally, we leveraged the BrainMap database to independently evaluate the neurobiological validity of the developed outrage signature. In line with previous work^46^, the thresholded MONS derived from the bootstrap test (FDR *q*<0.05; positive values only) was spatially overlapped with the modified 279-region version of the Brainnetome Atlas^75,76^. To quantify the spatial distribution of the MONS, we calculated the voxel contribution of each Brainnetome subregion, defined as the proportion of all MONS-positive voxels falling within that region. This analysis enabled us to identify the regions contributing most prominently to the spatial architecture of the outrage signature. The twenty regions showing the largest contributions were subsequently displayed. To further probe the functions of these regions, the functional profiles of twelve exemplary regions provided by the Brainnetome Atlas platform (https://atlas.brainnetome.org/bnatlas.html) were reported through probabilistic maps indicating the behavioral domain based on metadata taxonomy of the BrainMap database (http://www.brainmap.org/taxonomy/)^75^.

### Testing anger and disgust as components of outrage

Previous theoretical and behavioral accounts have suggested that outrage is a coupling of anger and disgust^2–5,13,50,51^. To examine whether this relationship is also evident on the neural level, we applied the MONS to independent datasets probing anger-related aggression and subjective disgust, respectively. First, we analyzed an independent TAP fMRI dataset (study 3), in which participants completed a well-established provocation-based aggression task probing state anger and reactive aggression (retaliatory punishment)^56,57^. The primary conditions of interest involved aggression following high versus low provocation. Participants were further stratified according to whether they exhibited increased post-task state anger, and we tested whether the MONS could differentiate aggression following high versus low provocation in the two subgroups (subgroup 1: *n*=41; subgroup 2: *n*=27; Supplementary Table 5) (see Supplementary Methods for experimental details). Second, we applied the MONS to two independent fMRI datasets^46^ (studies 4 and 5; *n*=78 and 30, respectively; Supplementary Table 5) in which participants viewed visual disgust stimuli eliciting varying levels of subjective disgust (for experimental design, see Supplementary Methods). Forced-choice classification was used to assess whether MONS responses discriminated between aggression following high versus low provocation across the two TAP subgroups and between different levels of subjective disgust. Pattern responses were calculated using the dot product of the MONS weight map with each participant’s brain activation map.

### Comparing signatures of outrage, negative affect and affective arousal

Building on prior conceptual and behavioral evidence suggesting both similarities and differences among outrage, negative affect and affective arousal^3,15,17^, we adopted a hypothesis-informed framework for comparing their neural signatures. Previous studies have developed and evaluated emotional signatures for general negative affect (MNAS^47^) and affective arousal (BAAS^48^). Like the MONS, the MNAS and BAAS used whole-brain (gray matter) voxel-wise activation maps (i.e., whole-brain beta maps derived from GLM analyses) as features. The MNAS model was developed to predict general negative affect induced by stimuli across modalities (visual, auditory and somatic) using partial least squares regression, with beta maps of each stimulus level as features and the ratings for each stimulus level as labels^47^. The BAAS model was developed to predict affective arousal evoked by videos using SVR, with beta maps from each rating as features and the ratings as labels^48^.

First, we examined the spatial similarities between stable decoding maps obtained from bootstrap tests and a priori cortical ROIs (dLPFC, vLPFC, dMPFC, vMPFC, OFC, ACC, MCC, PCC, anterior insula, middle insula, TPJ, and precuneus) as well as subcortical ROIs (PAG, hypothalamus, amygdala, thalamus, and ventral striatum) previously implicated in outrage, negative affect and arousal processes^9,47,48,61,62^ using river plots. Consistent with recent studies^47,103^, spatial similarity was computed as cosine similarity between the ROI and the thresholded MONS, MNAS and BAAS maps (FDR *q*<0.05, retaining positive values) from bootstrap tests.

Second, we examined the functional similarities between the MONS, MNAS and BAAS by applying the three signatures to the validation cohort (including both outrage- and punishment- related conditions), the MNAS multimodal negative affect dataset^47^ (study 6, *n*=55; Supplementary Methods and Supplementary Table 5) and the BAAS validation cohort^48^ (study 7, *n*=36; Supplementary Methods and Supplementary Table 5). Two-alternative forced-choice classification accuracies between different intensity levels of outrage (as well as punishment, negative affect and affective arousal) were calculated based on the respective pattern responses. Pattern responses were calculated using the dot product of each pattern weight map with the fMRI activation map for each participant.

Finally, we performed two-path mediation analyses^104^ using the Mediation Toolbox (https://github.com/canlab/MediationToolbox) to explore the relationship between MONS response, outrage rating, and MNAS response (for the principles of mediation analysis, see Supplementary Methods). Specifically, two multilevel mediation models were constructed: (1) the trial-by-trial MONS responses were entered as predictors (*X*), outrage ratings as outcomes (*Y*), and the trial-by-trial MNAS responses as mediators (*M*); (2) the trial-by-trial MNAS responses were entered as predictors (*X*), outrage ratings as outcomes (*Y*), and the trial-by-trial MONS responses as mediators (*M*). To do this, pattern responses for the MONS and MNAS were calculated for each participant as the dot product between single-trial beta maps and the corresponding signature, with trials showing VIFs greater than 3 excluded from analysis. Statistical significance of the mediation effects was assessed using bootstrap tests with 10,000 iterations. Mediation effects were considered significant when the bootstrapped 95% CI did not include zero (*P*<0.05). Additionally, we also tested whether (1) BAAS response mediated the relationship between MONS response and outrage ratings; (2) MONS response mediated the relationship between BAAS response and outrage ratings; (3) MNAS response mediated the relationship between MONS response and punishment ratings; (4) MONS response mediated the relationship between MNAS response and punishment ratings; (5) BAAS response mediated the relationship between MONS response and punishment ratings; and (6) MONS response mediated the relationship between BAAS response and punishment ratings.

### Validation in traditional moral judgment contexts

Prior work has demonstrated that outrage is closely associated with moral judgment^35,50,52,53^. To examine whether the outrage signature also tracks perceived moral wrongness, the MONS was applied to six independent moral judgment fMRI datasets obtained from ref.^54^, including four vignette-based datasets (studies 8–11; *n*=64, 30, 27 and 30, respectively; Supplementary Methods and Supplementary Table 5) and two picture-based datasets (studies 12–13; *n*=30 and 30, respectively; Supplementary Methods and Supplementary Table 5). Unlike the dynamic, audiovisual short-form videos used to elicit outrage in our study, these six datasets involved tasks that placed greater emphasis on explicit cognitive evaluation of moral violations, based on static images or written descriptions of hypothetical scenarios. In the vignette-based datasets, participants viewed vignettes of moral violations and rated the degree of moral wrongness using a 4-point Likert scale ranging from 1 (not morally wrong) to 4 (extremely morally wrong). Forced-choice classification accuracies were computed based on pattern responses to discriminate between low (level 1), moderate (average of levels 2 and 3) and high (level 4) levels of moral wrongness. In the picture-based datasets, participants rated the morality of images depicting moral violations on a 5-point bipolar Likert scale, ranging from 1 (very moral) to 5 (very immoral), with 3 indicating a neutral evaluation. Because this scale indexed morality and immorality rather than graded levels of moral wrongness, forced-choice classification analyses were restricted to comparisons between highly immoral (level 5) and neutral (level 3) stimuli, as well as between highly immoral (level 5) and highly moral (level 1) stimuli. Pattern responses were calculated as the dot product of the MONS weight map and each participant’s brain activation map. Given the differences in stimulus modality between the dynamic, naturalistic audiovisual short-form videos used to train the MONS and the static images or written vignettes used in the six validation datasets, we examined whether the MONS excluding the visual and auditory cortices could predict moral judgment across these datasets, to reduce the contribution of sensory processing associated with the training stimuli. We also evaluated the predictive performance of the whole-brain MONS for comparison.

## Acknowledgments

This work was partly supported by the China MOST2030 Brain Project (Grant Number: 2022ZD0208500) to D.Y., the RGC Humanities and Social Sciences Prestigious Fellowship Scheme (Grant Number: RGC 37000626) to B.B., RGC GRF (Grant Number: 17617526 and 17615525) to B.B., HKU seed grant for collaborative research (Grant Number: 2407102536) to B.B., start-up funding from The University of Hong Kong to B.B., the National Natural Science Foundation of China (Grant Number: 32601432) to X.G., and the Postdoctoral Fellowship Program of China Postdoctoral Science Foundation (Grant Number: GZC20261571) to X.G.

## References

1. Van Bavel, J. J., Robertson, C. E., Del Rosario, K., Rasmussen, J. & Rathje, S. Social media and morality. Annu. Rev. Psychol. 75, 311–340 (2024).

2. McLoughlin, K. L., Brady, W. J., Goolsbee, A., Kaiser, B., Klonick, K. & Crockett, M. J. Misinformation exploits outrage to spread online. Science 386, 991–996 (2024).

3. Brady, W. J., McLoughlin, K., Doan, T. N. & Crockett, M. J. How social learning amplifies moral outrage expression in online social networks. Sci. Adv. 7, eabe5641 (2021).

4. Brady, W. J., Crockett, M. J. & Van Bavel, J. J. The MAD model of moral contagion: the role of motivation, attention, and design in the spread of moralized content online. Perspect. Psychol. Sci. 15, 978–1010 (2020).

5. Brady, W. J., McLoughlin, K. L., Torres, M. P., Luo, K. F., Gendron, M. & Crockett, M. J. Overperception of moral outrage in online social networks inflates beliefs about intergroup hostility. *Nat*. Hum. Behav. 7, 917–927 (2023).

6. Crockett, M. J. Moral outrage in the digital age. *Nat*. Hum. Behav. 1, 769–771 (2017).

7. Spring, V. L., Cameron, C. D. & Cikara, M. The upside of outrage. Trends Cogn. Sci. 22, 1067–1069 (2018).

8. Robertson, C. E., Shariff, A. & Van Bavel, J. J. Morality in the anthropocene: the perversion of compassion and punishment in the online world. PNAS nexus 3, pgae193 (2024).

9. Gan, X., et al. Does unfairness evoke anger or disgust? A quantitative neurofunctional dissection based on 25 years of neuroimaging. Neurosci. Biobehav. Rev. 178, 106356 (2025).

10. Fehr, E. & Fischbacher, U. Third-party punishment and social norms. Evol. Hum. Behav. 25, 63–87 (2004).

11. Fehr, E. & Gächter, S. Altruistic punishment in humans. Nature 415, 137–140 (2002).

12. Boyd, R., Gintis, H., Bowles, S. & Richerson, P. J. The evolution of altruistic punishment. Proc. Natl. Acad. Sci. U. S. A. 100, 3531–3535 (2003).

13. Haidt, J. The moral emotions. In Handbook of affective sciences (eds Davidson, R. J., Scherer, K. R., Goldsmith, H. H.) (pp. 852–870). Oxford University Press (2003).

14. Brady, W. J., Jackson, J. C., Lindström, B. & Crockett, M. J. Algorithm-mediated social learning in online social networks. Trends Cogn. Sci. 27, 947–960 (2023).

15. Brady, W. J., Gantman, A. P. & Van Bavel, J. J. Attentional capture helps explain why moral and emotional content go viral. J. Exp. Psychol. Gen. 149, 746–756 (2020).

16. Gantman, A. P. & Van Bavel, J. J. Moral perception. Trends Cogn. Sci. 19, 631–633 (2015).

17. Rathje, S. & Van Bavel, J. J. The psychology of virality. Trends Cogn. Sci. 29, 914–927 (2025).

18. Robertson, C. E., Pröllochs, N., Schwarzenegger, K., Pärnamets, P., Van Bavel, J. J. & Feuerriegel, S. Negativity drives online news consumption. *Nat*. Hum. Behav. 7, 812–822 (2023).

19. Soroka, S., Fournier, P. & Nir, L. Cross-national evidence of a negativity bias in psychophysiological reactions to news. Proc. Natl. Acad. Sci. U. S. A. 116, 18888–18892 (2019).

20. Datareportal. Global social media Statistics, https://datareportal.com/social-media-users (accessed October 2026.).

21. Brady, W. J. & Crockett, M. J. Norm psychology in the digital age: how social media shapes the cultural evolution of normativity. Perspect. Psychol. Sci. 19, 62–64 (2024).

22. Robertson, C. E., Del Rosario, K. S. & Van Bavel, J. J. Inside the funhouse mirror factory: how social media distorts perceptions of norms. Current opinion in psychology 60, 101918 (2024).

23. Brady, W. J., et al. Redesigning algorithms to intervene on social norm misperceptions during a national election. Nature 655, 942–956 (2026).

24. Brady, W. J. & Crockett, M. J. How effective is online outrage? Trends Cogn. Sci. 23, 79–80 (2019).

25. Rathje, S., Van Bavel, J. J. & van der Linden, S. Out-group animosity drives engagement on social media. Proc. Natl. Acad. Sci. U. S. A. 118, e2024292118 (2021).

26. Williams, J. Stand out of our light: freedom and resistance in the attentional economy. Cambridge University Press (2018).

27. Gan, X., et al. Action real-time strategy gaming experience related to increased attentional resources: an attentional blink study. Front. Hum. Neurosci. 14, 101 (2020).

28. Milli, S., Carroll, M., Pandey, S., Wang, Y. & Dragan, A. D. Twitter’s algorithm: amplifying anger, animosity, and affective polarization. arXiv, (2023).

29. Chakradhar, S. More internal documents show how Facebook’s algorithm prioritized anger and posts that triggered it, https://www.niemanlab.org/2021/10/more-internal-documents-show-how-facebooks-algorithm-prioritized-anger-and-posts-that-triggered-it/ (assessed May 2026.). Nieman Lab (2021).

30. Spring, M. & Radford, M. Meta and TikTok let harmful content rise after evidence outrage drove engagement, say whistleblowers, https://www.bbc.com/news/articles/cqj9kgxqjwjo (accessed March 2026.). BBC (2026).

31. Lee, J. Y. The effects of social media criticism against public health institutions on trust, emotions, and social media engagement. Proc. Natl. Acad. Sci. U. S. A. 122, e2422890122 (2025).

32. Lee, J. Y. Reply to Loh: Evaluating anger as a mediator of criticism’s effect on social media engagement using the interventional effects approach. Proc. Natl. Acad. Sci. U. S. A. 122, e2529782122 (2025).

33. Lindström, B., Bellander, M., Schultner, D. T., Chang, A., Tobler, P. N. & Amodio, D. M. A computational reward learning account of social media engagement. Nat. Commun. 12, 1311 (2021).

34. Goldenberg, A. & Willer, R. Amplification of emotion on social media. *Nat*. Hum. Behav. 7, 845–846 (2023).

35. Brady, W. J., Wills, J. A., Jost, J. T., Tucker, J. A. & Van Bavel, J. J. Emotion shapes the diffusion of moralized content in social networks. Proc. Natl. Acad. Sci. U. S. A. 114, 7313–7318 (2017).

36. Oxford. The Oxford Word of the Year 2025 is rage bait, https://corp.oup.com/news/the-oxford-word-of-the-year-2025-is-rage-bait/ (accessed December 2025.). Oxford University Press (2025).

37. Hofmann, W., Wisneski, D. C., Brandt, M. J. & Skitka, L. J. Morality in everyday life. Science 345, 1340–1343 (2014).

38. Nguyen, L., et al. Feeds, feelings, and focus: a systematic review and meta-analysis examining the cognitive and mental health correlates of short-form video use. Psychol. Bull. 151, 1125–1146 (2025).

39. Montag, C., Yang, H. & Elhai, J. D. On the psychology of TikTok use: a first glimpse from empirical findings. Frontiers in public health 9, 641673 (2021).

40. Weaverdyck, M., Lieberman, M. & Parkinson, C. Multivoxel pattern analysis in fMRI: a practical introduction for social and affective neuroscientists. Soc. Cogn. Affect. Neurosci. 15, 487–509 (2020).

41. Kragel, P. A. & LaBar, K. S. Decoding the nature of emotion in the brain. Trends Cogn. Sci. 20, 444–455 (2016).

42. Kragel, P. A., Koban, L., Barrett, L. F. & Wager, T. D. Representation, pattern information, and brain signatures: from neurons to neuroimaging. Neuron 99, 257–273 (2018).

43. Haynes, J.-D. A primer on pattern-based approaches to fMRI: principles, pitfalls, and perspectives. Neuron 87, 257–270 (2015).

44. Woo, C.-W., Chang, L. J., Lindquist, M. A. & Wager, T. D. Building better biomarkers: brain models in translational neuroimaging. Nat. Neurosci. 20, 365–377 (2017).

45. Zhou, F., et al. A distributed fMRI-based signature for the subjective experience of fear. Nat. Commun. 12, 6643 (2021).

46. Gan, X., et al. A neurofunctional signature of subjective disgust generalizes to oral distaste and socio-moral contexts. *Nat*. Hum. Behav. 8, 1383–1402 (2024).

47. Čeko, M., Kragel, P. A., Woo, C.-W., López-Solà, M. & Wager, T. D. Common and stimulus-type-specific brain representations of negative affect. Nat. Neurosci. 25, 760–770 (2022).

48. Zhang, R., et al. A neurofunctional signature of affective arousal generalizes across valence domains and distinguishes subjective experience from autonomic reactivity. Nat. Commun. 16, 6492 (2025).

49. Kauvar, I., et al. Conserved brain-wide emergence of emotional response from sensory experience in humans and mice. Science 388, eadt3971 (2025).

50. Salerno, J. M. & Peter-Hagene, L. C. The interactive effect of anger and disgust on moral outrage and judgments. Psychol. Sci. 24, 2069–2078 (2013).

51. Brady, W. J., Wills, J. A., Burkart, D., Jost, J. T. & Van Bavel, J. J. An ideological asymmetry in the diffusion of moralized content on social media among political leaders. J. Exp. Psychol. Gen. 148, 1802–1813 (2019).

52. Rozin, P., Lowery, L., Imada, S. & Haidt, J. The CAD triad hypothesis: a mapping between three moral emotions (contempt, anger, disgust) and three moral codes (community, autonomy, divinity). J. Pers. Soc. Psychol. 76, 574–586 (1999).

53. Hutcherson, C. A. & Gross, J. J. The moral emotions: a social-functionalist account of anger, disgust, and contempt. J. Pers. Soc. Psychol. 100, 719–737 (2011).

54. Hopp, F., Youk, S., Sinnott-Armstrong, W. & Weber, R. A sensitive and specific neural signature robustly predicts graded computations of moral wrongness. Prepint, (2025).

55. Haufe, S., et al. On the interpretation of weight vectors of linear models in multivariate neuroimaging. Neuroimage 87, 96–110 (2014).

56. Akkoc Altinok, D. C., et al. A combined administration of testosterone and arginine vasopressin affects aggressive behavior in males. Brain Sci. 11, 1623 (2021).

57. Wagels, L., et al. Exogenous testosterone and the monoamine-oxidase A polymorphism influence anger, aggression and neural responses to provocation in males. Neuropharmacology 156, 107491 (2019).

58. Chang, L. J., Gianaros, P. J., Manuck, S. B., Krishnan, A. & Wager, T. D. A sensitive and specific neural signature for picture-induced negative affect. PLoS Biol. 13, e1002180 (2015).

59. Taschereau-Dumouchel, V., Kawato, M. & Lau, H. Multivoxel pattern analysis reveals dissociations between subjective fear and its physiological correlates. Mol. Psychiatry 25, 2342–2354 (2020).

60. Kohoutová, L., et al. Toward a unified framework for interpreting machine-learning models in neuroimaging. Nat. Protoc. 15, 1399–1435 (2020).

61. Blair, R. J., Habel, U., Votinov, M., Wagels, L. & Denson, T. F. A cognitive neuroscience approach to understanding aggression and its treatment. Neurosci. Biobehav. Rev. 179, 106386 (2025).

62. Alia-Klein, N., et al. The feeling of anger: from brain networks to linguistic expressions. Neurosci. Biobehav. Rev. 108, 480–497 (2020).

63. Amodio, D. M. & Frith, C. D. Meeting of minds: the medial frontal cortex and social cognition. Nat. Rev. Neurosci. 7, 268–277 (2006).

64. Mahmoodi, A. & Rushworth, M. F. S. Computational origins of cortical brain circuits for social cognition. Nat. Rev. Neurosci. 27, 345–356 (2026).

65. Miao, Z., et al. Common and distinct neural correlates of social interaction processing and theory of mind in narratives. Nat. Commun. 17, 4830 (2026).

66. Buergi, N., Aydogan, G., Konovalov, A. & Ruff, C. C. A neural signature of adaptive mentalization. Nat. Neurosci. 29, 934–944 (2026).

67. Berthoz, S., Armony, J. L., Blair, R. J. & Dolan, R. J. An fMRI study of intentional and unintentional (embarrassing) violations of social norms. Brain 125, 1696–1708 (2002).

68. Greene, J. D., Nystrom, L. E., Engell, A. D., Darley, J. M. & Cohen, J. D. The neural bases of cognitive conflict and control in moral judgment. Neuron 44, 389–400 (2004).

69. McAuliffe, K., Blake, P. R., Steinbeis, N. & Warneken, F. The developmental foundations of human fairness. *Nat*. Hum. Behav. 1, 0042 (2017).

70. Craig, A. D. How do you feel? Interoception: the sense of the physiological condition of the body. Nat. Rev. Neurosci. 3, 655–666 (2002).

71. Craig, A. D. How do you feel — now? The anterior insula and human awareness. Nat. Rev. Neurosci. 10, 59–70 (2009).

72. Andersen, R. A. Inferior parietal lobule function in spatial perception and visuomotor integration. In Comprehensive Physiology) (pp. 483-518). Wiley Online Library (2011).

73. Holmes, N. P., et al. Transcranial magnetic stimulation over supramarginal gyrus stimulates primary motor cortex directly and impairs manual dexterity: implications for TMS focality. J. Neurophysiol. 131, 360–378 (2024).

74. Gan, X., et al. Common and distinct neurofunctional representations of core and social disgust in the brain: coordinate-based and network meta-analyses. Neurosci. Biobehav. Rev. 135, 104553 (2022).

75. Fan, L., et al. The human brainnetome atlas: a new brain atlas based on connectional architecture. Cereb. Cortex 26, 3508–3526 (2016).

76. Lee, J.-J., et al. A neuroimaging biomarker for sustained experimental and clinical pain. Nat. Med. 27, 174–182 (2021).

77. Koban, L., Jepma, M., López-Solà, M. & Wager, T. D. Different brain networks mediate the effects of social and conditioned expectations on pain. Nat. Commun. 10, 4096 (2019).

78. Boyd, R. & Richerson, P. J. Punishment allows the evolution of cooperation (or anything else) in sizable groups. Ethology and Sociobiology 13, 171–195 (1992).

79. Fehr, E. & Rockenbach, B. Human altruism: economic, neural, and evolutionary perspectives. Curr. Opin. Neurobiol. 14, 784–790 (2004).

80. Spring, V. L., Cameron, C. D. & Cikara, M. Asking different questions about outrage: a reply to Brady and Crockett. Trends Cogn. Sci. 23, 80–82 (2019).

81. Zhu, J., Zhang, X., Liu, X. & Mu, Y. Neural architecture of social punishment: insights from a queue-jumping scenario. iScience 28, 111988 (2025).

82. Lindquist, K. A., Wager, T. D., Kober, H., Bliss-Moreau, E. & Barrett, L. F. The brain basis of emotion: a meta-analytic review. Behav. Brain Sci. 35, 121–143 (2012).

83. Clore, G. L. & Ortony, A. Appraisal theories: how cognition shapes affect into emotion. In Handbook of emotions, 3rd ed. (eds Lewis, M., Haviland-Jones, J. M., Barrett, L. F.) (pp. 628–642). The Guilford Press (2008).

84. Barrett, L. F. The theory of constructed emotion: an active inference account of interoception and categorization. Soc. Cogn. Affect. Neurosci. 12, 1–23 (2017).

85. Ridderinkhof, K. R., Ullsperger, M., Crone, E. A. & Nieuwenhuis, S. The role of the medial frontal cortex in cognitive control. Science 306, 443–447 (2004).

86. Dolan, R. J. The human amygdala and orbital prefrontal cortex in behavioural regulation. Philos. Trans. R. Soc. Lond. B. Biol. Sci. 362, 787–799 (2007).

87. Wallis, J. D. The dynamics of learning and behavioral flexibility. Neuron 71, 959–961 (2011).

88. Haber, S. N. & Knutson, B. The reward circuit: linking primate anatomy and human imaging. Neuropsychopharmacology 35, 4–26 (2010).

89. Knutson, B., Adams, C. M., Fong, G. W. & Hommer, D. Anticipation of increasing monetary reward selectively recruits nucleus accumbens. J. Neurosci. 21, Rc159 (2001).

90. Rathje, S., Robertson, C., Brady, W. J. & Van Bavel, J. J. People think that social media platforms do (but should not) amplify divisive content. Perspect. Psychol. Sci. 19, 781–795 (2024).

91. Molho, C., Tybur, J. M., Güler, E., Balliet, D. & Hofmann, W. Disgust and anger relate to different aggressive responses to moral violations. Psychol. Sci. 28, 609–619 (2017).

92. Molho, C., Tybur, J. M., Van Lange, P. A. M. & Balliet, D. Direct and indirect punishment of norm violations in daily life. Nat. Commun. 11, 3432 (2020).

93. Peirce, J. W. PsychoPy--Psychophysics software in Python. J. Neurosci. Methods 162, 8–13 (2007).

94. Jiang, H., et al. Common and distinct neurofunctional signatures of dynamic naturalistic emotion regulation strategies. Nat. Commun. 17, 4272 (2026).

95. Zhou, F., et al. Empathic pain evoked by sensory and emotional-communicative cues share common and process-specific neural representations. eLife 9, e56929 (2020).

96. Poldrack, R. A., Huckins, G. & Varoquaux, G. Establishment of best practices for evidence for prediction: a review. JAMA Psychiatry 77, 534–540 (2020).

97. Reddan, M. C., Wager, T. D. & Schiller, D. Attenuating neural threat expression with imagination. Neuron 100, 994–1005.e1004 (2018).

98. Atlas, L. Y., Lindquist, M. A., Bolger, N. & Wager, T. D. Brain mediators of the effects of noxious heat on pain. Pain 155, 1632–1648 (2014).

99. Büchel, C., Bornhovd, K., Quante, M., Glauche, V., Bromm, B. & Weiller, C. Dissociable neural responses related to pain intensity, stimulus intensity, and stimulus awareness within the anterior cingulate cortex: a parametric single-trial laser functional magnetic resonance imaging study. J. Neurosci. 22, 970–976 (2002).

100. Courville, T. & Thompson, B. Use of structure coefficients in published multiple regression articles: β is not enough. Educ. Psychol. Meas. 61, 229–248 (2001).

101. Thompson, B. & Borrello, G. M. The importance of structure coefficients in regression research. Educ. Psychol. Meas. 45, 203–209 (1985).

102. Yeo, B. T., et al. The organization of the human cerebral cortex estimated by intrinsic functional connectivity. J. Neurophysiol. 106, 1125–1165 (2011).

103. Gan, X., et al. Disgust propensity, not disgust sensitivity, shapes the reactivity of a subjective disgust circuit in humans. Hum. Brain Mapp. 47, e70577 (2026).

104. Woo, C.-W., et al. Quantifying cerebral contributions to pain beyond nociception. Nat. Commun. 8, 14211 (2017).

