## Supplementary Information for "Rage bait: a neurofunctional signature of moral outrage elicited by social media"

Gan et al.

Contact:
Dezhong Yao; Benjamin Becker

### Supplementary Methods

#### Analysis of trial-wise correspondence between outrage and punishment ratings

To characterize the trial-wise correspondence between outrage and punishment ratings within individuals, we constructed aligned rating profiles by sorting trials according to outrage ratings. Specifically, for each participant, the 60 trials were first ordered in ascending order based on outrage ratings. The corresponding punishment ratings were then rearranged using the same sorting indices, thereby preserving the trial-wise pairing between the two measures.

This procedure yielded, for each participant, a pair of aligned rating vectors in which outrage ratings increased monotonically across trials, allowing the corresponding pattern of punishment ratings to be directly compared. Group-level mean trajectories and standard errors of the mean (s.e.m.) were subsequently computed across participants for both outrage and punishment ratings at each sorted trial position.

#### MRI data acquisition for the discovery and validation cohorts

High-resolution structural images were obtained with a T1-weighted spoiled gradient recalled (SPGR) sequence (repetition time (TR)=6 ms, echo time (TE)=2 ms, flip angle=9°, field of view (FOV)=256 × 256 $mm^2$, voxel size=1 × 1 × 1 $mm^3$, acquisition matrix=256 × 256, 156 slices, slice thickness=1 mm). These structural images were used to facilitate spatial normalization and to screen for potential brain abnormalities. Functional images were acquired using a T2*-weighted echo-planar imaging (EPI) sequence (TR=2000 ms, TE=30 ms, flip angle=90°, FOV=240 × 240 $mm^2$, voxel size=3.75 × 3.75 × 4 $mm^3$, matrix size=64 × 64, 39 slices, slice thickness=3 mm with a 1 mm inter-slice gap).

#### First-level fMRI analysis of punishment in the validation cohort

In this general linear model (GLM), neural responses during the clip presentation period were modeled using five condition-specific regressors corresponding to the binned punishment ratings (levels 1–5), enabling separate estimation of brain activity associated with each rating level. In addition, two boxcar regressors (4 s) were included to capture activity during the two rating periods (i.e., outrage and punishment rating phases, respectively), thereby accounting for motor-related processes. Fixation periods were treated as the implicit baseline. All task-related regressors were convolved with the canonical hemodynamic response function. Functional runs were concatenated within each participant using the SPM function spm_fmri_concatenate.m. Nuisance regressors included: (1) run-specific intercepts to account for mean signal differences across runs; (2) head motion parameters (three translations and three rotations), along with their squares, derivatives and squared derivatives (24 regressors in total); and (3) regressors indexing time points identified as motion-related outliers.

#### Determining brain regions associated with and predictive of subjective outrage on the individual level

To further characterize neural circuits involved in the subjective experience of outrage on the individual level, we combined complementary univariate and multivariate analytic approaches. First, brain regions showing significant associations with subjective outrage ratings were identified using the one-sample *t* test on the first-level univariate parametric modulation beta maps derived from the GLM analyses. We next employed multivariate analyses to localize regions that were predictive of and associated with outrage ratings separately, as well as brain regions showing an overlapping effect. Specifically, we conducted within-subject multivariate analyses to determine brain regions that reliably contributed to the prediction of outrage ratings across individuals. Briefly, for each participant, a prediction model (linear SVR; $C$=1) was trained on single-trial activation maps using a 10×10-fold cross-validation procedure. The consistency of voxel-wise weights obtained from these within-subject multivariate models was evaluated using a one-sample $t$ test. Regions surviving FDR correction ($q$<0.05) were considered to exhibit consistent outrage-predictive effects across participants. We only included participants whose trial-wise outrage ratings could be significantly predicted by their brain data (assessed using prediction–outcome Pearson correlation; $n$=71, from 80 participants). Importantly, comparable findings were obtained when all participants were included. Moreover, within-subject predictive patterns were transformed into 'activation patterns' to identify regions showing consistent outrage-related effects across individuals (thresholded at $q$<0.05, FDR corrected). Finally, as with the population-level model, we identified the overlap between the within-subject multivariate patterns and 'activation patterns', which reflected regions that were both significantly predictive of and associated with the subjective experience of outrage on the individual level.

**Modified Taylor aggression paradigm (TAP) used in study 3**

The TAP used in study 3[1,2] was implemented as a competitive reaction time game task in which participants competed against an ostensible opponent. Participants were instructed to press a button as quickly as possible when a soccer ball appeared on the screen. They were informed that they were competing against a real participant (although the opponent's behavior was in fact computer-controlled), and that the opponent would subtract some money from them when they responded more slowly than the opponent. Each trial lasted 15–16 s, and the entire task duration was 25 min.

At the start of each trial, participants selected the amount of money (0–100 cents) they wished to subtract from the opponent in case the opponent lost. Responses were made using a visual analogue scale. This retaliatory decision was interpreted as an index of reactive aggression in response to the provocation (i.e., money subtracted by the opponent) experienced during the preceding trial. Participants were explicitly informed that deducted money would not be transferred to themselves, ensuring that the subtraction behavior reflected intentional punishment toward the provoking opponent rather than personal monetary gain. During the subsequent reaction-time phase, participants were instructed to respond as rapidly as possible to the appearance of the moving soccer ball. At the end of each trial, participants received feedback indicating whether they had won or lost and, following loss trials, the amount of money deducted by the ostensible opponent. Loss trials therefore served as provocation conditions, whereas win trials, during which no opponent decision was displayed, served as non-provocation control conditions. There were 54 predefined loss trials, including 23 high-provocation trials (80–100 cents subtracted), 25 low-provocation trials (0–20 cents subtracted), and 6 medium-provocation trials (30–70 cents subtracted), as well as approximately 30 win trials in which participants always gained 50 cents. State anger ratings were additionally collected before and after completion of the TAP.

The primary conditions of interest were aggression following high provocation and low provocation in the placebo group. For the following analyses, the beta images of aggression after high and after low provocation were used. To further examine the relationship between the neural representation of outrage and provocation-induced anger experience, the placebo group was subdivided according to changes in state anger following the TAP. Specifically, one subgroup exhibited increased post-task state anger, whereas the other did not show an increase in state anger. In both subgroups, we examined whether the MONS could differentiate aggression following high versus low provocation.

**Subjective disgust rating fMRI paradigm used in studies 4–5**

Participants in study 4 and study 5 underwent the same disgust induction task[3], which incorporated 80 pictorial stimuli from the DIsgust-RelaTed-Images (DIRTI) disgust images database[4] (67 pictures), Nencki Affective Picture System (NAPS)[5] (5 pictures) and International Affective Picture System (IAPS)[6] (8 pictures), distributed over 4 runs (with 20 stimuli per run). Participants were instructed to pay attention to the pictures and naturally experience the induced emotion. Each trial consisted of a 6-s presentation of the picture followed by a 2-s fixation cross separating the stimuli from the rating period. Participants then had 4 s to report the level of disgust they experienced for the stimuli using a 5-point Likert scale, where 1 indicated neutral/slightest disgust and 5 indicated strongest disgust.

For each fMRI dataset, we tested whether the MONS could predict high (ratings 4–5) versus low (rating 1) disgust, high versus moderate (ratings 2–3) disgust, and moderate versus low disgust.

**Multimodal aversive experience task used in study 6**

In study 6[7], all participants were exposed to four intensity levels for each type of aversive stimulus (painful heat, painful pressure, aversive sounds and aversive images). Stimulus intensity was determined by factors such as temperature in heat pain studies or normative ratings in picture-based paradigms. After each stimulus, participants rated their negative affect ('aversiveness') in terms of avoidance ('how much do you want to avoid this experience in the future?'). In total, participants received 96 aversive stimuli (4 stimulus types × 4 intensities × 6 runs presented in random order).

For the comparisons of functional similarities in the present study, we defined low negative affect as rating 1, moderate negative affect as the average of ratings 2 and 3, and high negative affect as rating 4.

**Affective arousal rating fMRI paradigm used in study 7**

In study 7[8], participants were presented with 28 one-minute videos. These videos represented two affective states (9 positive clips and 9 negative clips) as well as a neutral state (10 neutral clips). During fMRI, all stimuli were presented in three runs, with either 3 or 4 clips from each category per run. After each video, participants rated their level of arousal during each video using a 9-point Likert scale, where 1 indicates very low arousal and 9 indicates very high arousal.

For the comparisons of functional similarities in the present study, we defined low affective arousal as the average of ratings 1–3, moderate affective arousal as the average of ratings 4–6, and high affective arousal as the average of ratings 7–9.

**Principles of mediation analysis**

Mediation analysis was used to examine whether the covariance between an independent (predictor) variable (*X*) and a dependent (outcome) variable (*Y*) could be accounted for by a third variable (*M*; mediator). The associations between predictor and mediator, mediator and outcome, as well as predictor and outcome before and after controlling for the mediator, are represented by paths *a*, *b*, *c*, and *c′*, respectively. Accordingly, the total effect of the predictor on the outcome (path *c*) can be decomposed into a direct effect (path *c′*) and an indirect (mediation) effect, quantified as the product of path coefficients *a* and *b* (that is, $a \times b$). A significant mediation effect was inferred when paths a and b, as well as the indirect effect ($a \times b$), reached statistical significance. Moreover, when path *c′* remained significant after accounting for the mediator, the mediation was interpreted as partial; otherwise, the mediator was considered to fully mediate the relationship between the predictor and outcome variables.

**Evaluating the specificity of MONS with arousal controlled**

To further determine the extent to which affective arousal contributes to the predictive performance of the MONS, we evaluated whether the signature retained its ability to discriminate subjective outrage after controlling for BAAS-derived arousal-related variance, adopting an approach conceptually similar to that employed in a recent study[8]. Pattern responses were quantified as the dot product between the MONS/BAAS and the corresponding brain activation map. First, the MONS was applied to the discovery (10×10-fold cross-validation) and independent validation cohorts to quantify pattern responses and evaluate classification performance across successive pairs of outrage intensity levels as well as low, moderate and high outrage. In the discovery cohort, pattern responses were averaged across the 10 repeated cross-validation iterations before statistical evaluation. Second, the BAAS response for each outrage intensity level in both the discovery and validation cohorts was computed. Finally, to assess whether MONS retained predictive utility beyond arousal-related processes, BAAS responses were regressed out from the corresponding MONS responses, and the adjusted MONS responses were subsequently used to re-evaluate discrimination performance across outrage intensity levels in both cohorts.

**Vignette-based moral judgment paradigm used in studies 8–11**

Studies 8–11 involved four vignette-based moral judgment datasets from ref.[9], corresponding to the moral vignettes discovery, validation, replication and generalization cohorts. During fMRI scanning, participants in the discovery and replication cohorts viewed the original Moral Foundations Vignettes[10] (MFV), whereas those in the generalization cohort viewed the validated Dutch version of the MFV[11]. One example vignette is *"You see a man leaving his family business to go work for their main competitor"*. The MFV comprise 120 brief descriptions of moral transgressions, each consisting of a single sentence (14–17 words) and targeting one of seven moral foundations: physical care, emotional care, fairness, liberty, loyalty, authority or sanctity. The stimulus set also includes a category of non-moral social norm violations. The validation cohort completed judgments for 72 vignettes. Of these, 42 were selected from the MFV, with six items representing each of the seven moral foundations. The remaining 30 vignettes were developed by the original authors to cover six moral domains specified in the Morality as Cooperation (MAC) framework[12]: family, reciprocity, bravery, property, respect, and group values, with five items per domain.

The vignettes were presented in an event-related fMRI design across three functional runs, each lasting approximately 8 min, with presentation order randomized. On each trial, participants read a vignette and were instructed to vividly imagine the situation it described. They then judged the moral wrongness of the depicted action using an MRI-compatible button box, with ratings ranging from 1 (not morally wrong) to 4 (extremely morally wrong). Participants had approximately 8 s to read and make judgments of each vignette.

#### Picture-based moral judgment paradigm used in studies 12–13

Studies 12–13 involved two picture-based moral judgment datasets from ref.[9]. In study 12, participants viewed 78 pictures, whereas study 13 included 120 pictures, 59 of which overlapped with those used in study 12. The pictures were representatively sampled from the Socio-Moral Image Database (SMID)[13], as described in ref.[11]. The sociomoral pictures are available at https://osf.io/2rqad/.

In each study, stimuli were presented across three functional runs, with 26 pictures per run in study 12 and 40 pictures per run in study 13. Each trial began with a 2-s fixation cross, followed by a 7-s picture presentation and a 4-s rating period. Participants were instructed to attend to each picture and judge whether the picture displays something moral or immoral using a 5-point bipolar Likert scale, ranging from 1 (very moral) to 5 (very immoral), with 3 indicating neutral. Ratings of 2 and 4 corresponded to somewhat moral and somewhat immoral, respectively.

#### Development of the moral judgment signature (MJS)

Like the MONS, the MJS used whole-brain (gray matter) voxel-wise activation maps (i.e., whole-brain beta maps derived from GLM analyses) as features. The MJS model was developed to predict moral wrongness ratings evoked by moral vignettes using SVR, with beta maps from each rating as features and the ratings as labels. For further details, please refer to ref.[9].

#### Examination of the spatial relationship between MONS and MJS

First, we assessed the whole-brain spatial correlation between MONS and MJS and evaluated its significance using permutation testing with 5,000 random shuffles. Next, to illustrate the voxel-level spatial similarity between MONS and MJS, we examined the joint distribution of normalized ($z$-scored) whole-brain voxel weights of these two signatures by plotting MONS on the $y$-axis and the MJS on the $x$-axis (for a similar approach, see ref.[14]). We then divided the resulting scatter plot into eight octants that coarsely define directions of shared positive or shared negative weights (Octant 2 and Octant 6, respectively), and uniquely MJS- or MONS-positive and -negative weights (unique MONS positive in Octant 1, negative in Octant 5; unique MJS positive in Octant 3, negative in Octant 7). Opposite weights for the two neural signatures would be seen in Octants 4 and 8.

### Supplementary Results

#### Distribution of binned outrage and punishment ratings across cohorts

Following the binning procedure, all participants in the discovery cohort used level 1 in their responses, whereas 73%, 98%, 98%, and 95% used levels 2–5, respectively, for outrage ratings; the corresponding proportions for punishment ratings (1–5) were 100%, 78%, 100%, 100%, and 100%. A similar pattern was observed in the validation cohort, where 100%, 77%, 97%, 93%, and 90% of participants used levels 1–5 for outrage, and 100%, 80%, 100%, 100%, and 93% for punishment.

#### Sensitivity of the MONS to different levels of outrage and punishment tendencies

To further examine whether the MONS could discriminate different levels of subjective outrage experience and punishment tendencies, we performed two-alternative forced-choice classification analyses based on pairs of activation maps within each participant. For each comparison, the activation map with the higher MONS response was classified as reflecting higher levels of outrage or punishment tendency, and classification accuracies were assessed against chance level (50%) using two-sided independent binomial tests.

For outrage ratings, the MONS reliably distinguished high (average of ratings 4–5) versus moderate (average of ratings 2–3) and moderate versus low (rating 1) levels in both discovery and validation cohorts, achieving accuracies of 90–96% (Cohen's $d$=1.95–2.89). High versus low outrage levels were classified with 100% accuracy in both cohorts (Cohen's $d$=4.76–4.96). The MONS also significantly discriminated adjacent outrage rating levels (e.g., rating 1 versus rating 2), with accuracies ranging from 77% to 93% ($P$<0.009) (Fig. 2b,c).

For punishment tendencies, the MONS showed comparable sensitivity. High versus moderate and moderate versus low punishment levels were classified with accuracies of 90–93% (Cohen's $d$=1.89–2.15), whereas high versus low punishment levels were classified with 100% accuracy ($P$<0.001, Cohen's $d$=4.60). Adjacent punishment levels were also significantly distinguished, with accuracies exceeding 82% ($P$<0.001), except for the comparison between rating levels 3 and 4 (accuracy=57%, $P$=0.58).

#### MONS prediction is largely preserved after excluding visual and auditory cortices

To assess the contribution of visual and auditory cortices to MONS performance, we retrained the outrage signature after excluding these sensory regions (the auditory cortex composed of A1, A2, and A3 from the HCP-MMP1.0 atlas[15]; the visual cortex composed of V1, V2, V3, and V4 from the HCP-MMP1.0 atlas[15]). The resulting signature largely maintained its predictive performance across cohorts and outcome measures. Specifically, the overall prediction–outcome correlations were $r$=0.53 (95% CI=[0.47, 0.61]) in the discovery cohort (cross-validation), $r$=0.47 (95% CI=[0.34, 0.58]) in the validation cohort when outrage was the outcome, and $r$=0.45 (95% CI=[0.32, 0.57]) in the validation cohort when punishment was the outcome. Mean within-subject correlations between predicted and true ratings were also maintained (discovery cohort, $r$=0.90±0.01; validation cohort, outrage as outcome, $r$=0.89±0.02; validation cohort, punishment as outcome, $r$=0.84±0.03). Classification accuracies were similarly preserved (Supplementary Fig. 2a,b,c). These findings suggest that the visual and auditory cortices are not essential for maintaining the predictive performance of the outrage signature, although they may still contribute to the neural representation of outrage.

### Within-subject trial-wise prediction using original ratings

To further evaluate the within-subject predictive performance of the MONS without grouping rating levels, we conducted additional prediction analyses using the original ratings (1–9) as trial-wise outcomes.

In the discovery cohort, the MONS trained using the five-level rating scheme significantly predicted trial-by-trial outrage ratings in 79 of 80 participants (cross-validated), yielding a mean prediction–outcome correlation of $r$=0.59±0.01 (mean±s.e.m., Supplementary Fig. 3a). Similar effects were observed in the independent validation cohort, where the MONS significantly predicted original outrage ratings in 29 of 30 participants (mean $r$=0.53±0.01; Supplementary Fig. 3b). The MONS also significantly predicted trial-wise punishment ratings in all validation participants ($n$=30; mean $r$=0.53±0.01; Supplementary Fig. 3c). Together, these findings show that the MONS trained using grouped ratings can capture trial-by-trial variations in subjective outrage and related punishment responses when evaluated against the original, ungrouped ratings, providing further support for the rating-grouping approach adopted in the primary analyses.

### Brain regions associated with and predictive of subjective outrage on the individual level

To further delineate regions underlying the subjective experience of outrage on the individual level, we combined convergent univariate and multivariate analyses. We first conducted a one-sample $t$ test on the first-level parametric modulation beta maps to determine brain regions that increased activity in association with the subjective experience of outrage (i.e., the higher the outrage ratings, the higher functional activation in the respective brain regions). Results revealed that subjective outrage was correlated with activation in a broad network spanning multiple cortical and subcortical systems, with increasing activation in a broadly distributed bilateral network encompassing the precuneus, dMPFC, dLPFC, vLPFC, ACC, MCC, PCC, middle frontal gyrus, inferior frontal gyrus, anterior and mid- and posterior insular cortex, superior temporal sulcus, superior and middle and inferior temporal gyrus, temporal pole, TPJ, fusiform gyrus, cuneus, supplementary motor area, precentral gyrus, inferior parietal lobule, supramarginal gyrus, hippocampus, thalamus, caudate, putamen, substantia nigra, hypothalamus, PAG, brainstem and cerebellar regions. In contrast, negative associations with outrage ratings were observed in the OFC, amygdala, and lingual gyrus (thresholded at FDR $q$<0.05 corrected for multiple comparisons; Supplementary Fig. 4a).

We next validated and extended the univariate results by using multivariate models in several ways. A one-sample $t$ test (treating participant as a random effect) was performed on the weights obtained from within-subject multivariate predictive models (for details, see Supplementary Methods). Similar to the univariate maps, within-subject predictive models (backward models) included consistent weights in brain regions spanning multiple large-scale cortical and subcortical systems that largely overlapped with the outrage regions as determined by the univariate parametric modulation approach (Supplementary Fig. 4b; $q$<0.05, FDR corrected). To further provide a more direct comparison between univariate and multivariate results, we transformed the within-subject outrage-predictive patterns into 'activation patterns' (or 'structure coefficients'), which map each voxel to the overall multivariate model prediction[16]. The resulting thresholded 'model activation patterns' (Supplementary Fig. 4c) exhibited marked overlap with both the univariate parametric modulation maps and the multivariate predictive-weight patterns (Supplementary Fig. 4a,b), indicating that the subjective experience of outrage is encoded in a distributed neural representation encompassing cortical and subcortical regions.

The overlap (i.e., conjunction) of within-subject model weights and model encoding maps (Supplementary Fig. 4d) revealed that most significant model weights in the multivariate models also encoded model information (i.e., showed significant 'model activation patterns'). Collectively, these findings indicate that the neural representation of human moral outrage is not limited to isolated focal regions (e.g., the thalamus), but is instead encoded in a broad set of regions spanning cortical and subcortical systems.

**MONS outperforms prediction based on local systems**

As previous studies emphasized the contribution of specific brain systems, such as mentalizing regions (e.g., precuneus, PCC)[17], threat detection regions (e.g., insula, thalamus, amygdala, hypothalamus, PAG)[17-19] and salience and default mode networks[19-21], to the subjective experience of outrage, we systematically tested whether local systems (searchlight-, parcellation-, ROI- and network-based) were sufficient to predict subjective outrage relative to the whole-brain MONS. As shown in Supplementary Fig. 5a,b, subjective outrage could be significantly predicted by activations in widely distributed regions ($P<0.001$, averaged across 10×10-fold cross-validation procedure). However, none of the local models predicted subjective outrage to the extent that the MONS did (see Supplementary Fig. 6a,b for predictions of models trained on the discovery cohort on the validation cohort).

We next re-trained predictive SVR models restricted to activations in (1) a set of cortical ROIs (i.e., the OFC, ACC, MCC, PCC, anterior insula, middle insula, TPJ and precuneus); (2) a set of subcortical ROIs (i.e., the PAG, hypothalamus, amygdala and thalamus); (3) a subcortical network; (4) each of seven large-scale cerebral networks (see Methods). Results showed that the cortical ROIs (prediction–outcome correlation $r$=[0.17, 0.32] and [0.14, 0.35] for discovery (10×10-fold cross-validation) and validation cohorts, respectively), the subcortical ROIs (prediction–outcome correlation $r$=[0.21, 0.30] and [0.15, 0.39] for discovery (10×10-fold cross-validation) and validation cohorts, respectively) as well as large-scale brain networks (Supplementary Fig. 5c,d,e, Supplementary Fig. 6c,d,e, and Supplementary Tables 3,4) could, to some extent, predict subjective outrage. Nonetheless, although statistically significant (except for the OFC, middle insula and thalamus in the validation cohort with $P>0.05$), the effect sizes in terms of prediction–outcome correlations (including searchlight- and parcellation-based predictions) were substantially smaller than those obtained from the MONS, which incorporated features distributed across multiple brain systems.

To account for potential influences of the number of features on prediction performance (that is, the substantially larger number of voxels included in the whole-brain model), we performed additional analyses based on random voxel sampling. Specifically, voxels were randomly sampled 1,000 times from a uniform contribution across the whole-brain (black), subcortical network (pink), or each large-scale cerebral network (other colored lines), and prediction performance was averaged across iterations[3,22,23]. As illustrated in Supplementary Fig. 5e and Supplementary Fig. 6e, the model constructed from voxels sampled across the entire brain (i.e., the MONS) achieved markedly higher asymptotic prediction performance (as indicated by much larger effect sizes) than models restricted to individual networks (colored lines; see also Supplementary Tables 3 and 4 for details). Importantly, prediction performance approached an asymptote when approximately 10,000 voxels were sampled across the whole-brain, indicating that robust prediction of subjective outrage relies on information distributed across multiple neural systems rather than confined to isolated networks. Comparable results were observed when models trained in the discovery cohort were applied to the independent validation cohort (Supplementary Fig. 6e), further demonstrating the robustness of prediction models trained on approximately 10,000 randomly sampled voxels (for similar observations, see refs.[3,8,22]). Together, these analyses consistently indicate that subjective outrage is encoded in distributed neural patterns spanning multiple cortical and subcortical systems, rather than being reducible to isolated regions or single large-scale networks. These findings further support the emerging view that emotions are represented in distributed brain-wide systems.

**Regional contributions of MONS, MNAS and BAAS**

We investigated spatial similarities between stable decoding maps and a set of cortical and subcortical ROIs (Supplementary Table 7). As shown in Fig. 6a,b, the PCC exhibited a preferential contribution to the MONS. The ACC, MCC, precuneus and anterior insula contained stable predictive voxels in both the MONS and MNAS, but the first three ROIs demonstrated relatively larger contributions to the MONS. The hypothalamus, PAG, thalamus, dMPFC, middle insula, TPJ, vLPFC, dLFPC and ventral striatum displayed reliable predictive contributions across all three models. Among these regions, the hypothalamus showed the largest contribution to the MONS, whereas the PAG contributed more strongly to the MONS and BAAS than to the MNAS. The thalamus and dMPFC contributed comparably to the three models. In contrast, although the middle insula and TPJ also exhibited strong involvement in the MONS, both ROIs showed relatively greater contributions to the BAAS and MNAS. The vLPFC, dLPFC and ventral striatum were largely specific to the BAAS and MNAS. Finally, the vMPFC, OFC and amygdala showed no stable positive contribution to the MONS. This finding for the OFC and amygdala should be interpreted with caution, however, because the spatial similarity analyses considered only positive model weights, whereas both regions displayed negative weights in the MONS (Fig. 2a). Rather than indicating that these regions are uninvolved in outrage, their contributions may instead be expressed through relative deactivation or opponent coding within the distributed representation of outrage.

**MONS response mediates the association between subjective outrage and neural reactivity of a multimodal general negative affect signature (MNAS)**

We performed multilevel mediation analyses to examine the relationship between MONS response, outrage ratings and MNAS response. We first tested whether the MONS response (*M*) could mediate the relationship between the MNAS response (*X*) and subjective outrage rating (*Y*). The results showed that, in the discovery cohort, the MNAS response was positively associated with the MONS response (path *a*; β=2.60, s.e.m.=0.56, 95% CI=[1.51, 3.70]; $Z$=3.74, $P$<0.001; Cohen's $d$=0.12), the MONS response remained positively associated with outrage ratings after controlling for the MNAS response (path *b*; β=0.38, s.e.m.=0.01, 95% CI=[0.35, 0.40]; $Z$=3.60, $P$<0.001; Cohen's $d$=0.75). In addition, there was a significant mediation effect (path $a \times b$; β=0.86, s.e.m.=0.18, 95% CI=[0.50, 1.21]; $Z$=3.68, $P$<0.001; Cohen's $d$=0.10). Furthermore, both the total effect (path *c*; β=1.64, s.e.m.=0.27, 95% CI=[1.12, 2.15]; $Z$=3.74, $P$<0.001; Cohen's $d$=0.12) and the non-mediated effect (path *c′*; β=0.72, s.e.m.=0.22, 95% CI=[0.29, 1.14]; $Z$=3.34, $P$=0.001; Cohen's $d$=0.07) were significant. Comparable findings were observed in the validation cohort, although path *a* only showed a trend toward significance ($P$=0.06). Together, these results indicate that the MONS response partially mediates the relationship between MNAS response and subjective outrage rating in both the discovery and validation cohorts (Fig. 6e).

Additionally, a second multilevel mediation model was constructed with the MONS response as *X*, the MNAS response as the mediator (*M*) and subjective outrage rating as *Y*. The results revealed that the MNAS did not significantly mediate the association between the MONS response and outrage rating in either the discovery or validation cohort (Fig. 6f), consistent with the relatively weak overall prediction–outcome correlations obtained when applying the MNAS to predict subjective outrage in the two cohorts (discovery cohort: $r$=0.12, 95% CI=[0.02, 0.21]; validation cohort: $r$=0.09, 95% CI=[-0.07, 0.25]), suggesting that the MONS captures neural representations that are, to a certain extent, more specific to outrage than general negative affect.

**MONS response during outrage is partially mediated by the affective arousal signature (BAAS)**

BAAS significantly predicted subjective outrage (discovery cohort: $r$=0.28, 95% CI=[0.18, 0.37]; validation cohort: $r$=0.24, 95% CI=[0.08, 0.39]), we thus further employed two multilevel mediation analyses to investigate the relationship between MONS response, BAAS response and subjective outrage rating. Specifically, we tested whether (1) the MONS response (*M*) could explain the relationship between the BAAS response (*X*) and outrage rating (*Y*), and whether (2) the BAAS response (*M*) could mediate the association between MONS response (*X*) and outrage rating (*Y*). The results showed that in the discovery cohort, the BAAS response was positively associated with the MONS response (path *a*; β=7.34, s.e.m.=0.45, 95% CI=[6.44, 8.24]; $Z$=3.65, $P$<0.001; Cohen's $d$=0.47), the MONS response was positively correlated with outrage rating (path *b*; β=0.35, s.e.m.=0.01, 95% CI=[0.32, 0.37]; $Z$=3.67, $P$<0.001; Cohen's $d$=0.64) and a significant mediation effect was observed (path $a \times b$; β=2.31, s.e.m.=0.13, 95% CI=[2.05, 2.57]; $Z$=3.58, $P$<0.001; Cohen's $d$=0.32). Moreover, both path *c* (β=3.85, s.e.m.=0.23, 95% CI=[3.40, 4.31]; $Z$=3.73, $P$<0.001; Cohen's $d$=0.39) and path *c′* (β=1.35, s.e.m.=0.21, 95% CI=[0.95, 1.79]; $Z$=3.86, $P$<0.001; Cohen's $d$=0.15) were significant. Overall, these results indicate that the MONS response partially mediated the effect of the BAAS response on the outrage rating. Notably, these findings were replicated in the validation cohort (Supplementary Fig. 8a).

We next constructed an alternative mediation model with BAAS response as *M*, MONS response as *X* and outrage rating as *Y*. Although a partial mediation effect of the BAAS response was observed, the former model (Supplementary Fig. 8a) showed larger effect sizes than the latter model (Supplementary Fig. 8b), with the mediation effect being 3.56 times greater in the discovery cohort (model 1: Cohen's $d$ = 0.32; model 2: Cohen's $d$ = 0.09). A similar trend was observed in the validation cohort. Taken together, these findings suggest that although affective arousal partially contributes to outrage-related neural responses, the MONS captures variance more directly linked to subjective outrage experience (for more evidence, see Supplementary Fig. 11).

**MONS response mediates the association between the MNAS and punishment rating**

We performed additional multilevel mediation analyses to explore the relationship between MONS response, punishment ratings and MNAS response. We first tested whether the MONS response (*M*) could mediate the relationship between the MNAS response (*X*) and punishment rating (*Y*). The results showed that, in the discovery cohort, the MNAS response was positively associated with the MONS response (path *a*; β=2.60, s.e.m.=0.56, 95% CI=[1.54, 3.74]; $Z$=3.83, $P$<0.001;

Cohen's $d$=0.12), while the MONS response remained positively associated with punishment ratings after controlling for the MNAS response (path $b$; β=0.39, s.e.m.=0.01, 95% CI=[0.37, 0.42]; $Z$=3.65, $P$<0.001; Cohen's $d$=0.74). There was a significant mediation effect (path $a \times b$; β=0.90, s.e.m.=0.19, 95% CI=[0.53, 1.29]; $Z$=3.77, $P$<0.001; Cohen's $d$=0.10). Furthermore, both the total effect (path $c$; β=1.50, s.e.m.=0.28, 95% CI=[0.96, 2.06]; $Z$=3.80, $P$<0.001; Cohen's $d$=0.10) and the direct effect (path $c'$; β=0.55, s.e.m.=0.23, 95% CI=[0.11, 1.00]; $Z$=2.42, $P$=0.01; Cohen's $d$=0.05) were significant. A similar mediation profile was observed in the validation cohort. Together, these results indicate that the MONS response plays a partial mediation role in the relationship between MNAS response and punishment rating in both the discovery and validation cohorts (Supplementary Fig. 9a).

A second multilevel mediation model was constructed with the MONS response as $X$, the MNAS response as the mediator ($M$) and punishment rating as $Y$. The results revealed that the MNAS failed to mediate the correlation between the MONS response and punishment rating in either the discovery or validation cohort (Supplementary Fig. 9b), which dovetailed with the relatively low overall prediction–outcome correlations obtained when applying the MNAS to predict punishment in the two cohorts (discovery cohort: $r$=0.08, 95% CI=[-0.02, 0.17]; validation cohort: $r$=0.06, 95% CI=[-0.09, 0.21]). These findings suggest that the MONS response to punishment tendency is not merely driven by neural representations related to general negative affect.

**MONS response toward punishment is partially mediated by the affective arousal signature (BAAS)**

BAAS could significantly predict punishment ratings (discovery cohort: $r$=0.25, 95% CI=[0.15, 0.34]; validation cohort: $r$=0.23, 95% CI=[0.07, 0.37]). We therefore conducted two additional multilevel mediation analyses to characterize the relationship between MONS response, BAAS response and punishment rating. Specifically, we examined whether (1) the MONS response ($M$) mediated the relationship between the BAAS response ($X$) and punishment rating ($Y$), and whether (2) the BAAS response ($M$) mediated the association between MONS response ($X$) and punishment rating ($Y$). The results demonstrated that, in the discovery cohort, the BAAS response was positively associated with the MONS response (path $a$; β=7.34, s.e.m.=0.46, 95% CI=[6.42, 8.20]; $Z$=3.60, $P$<0.001; Cohen's $d$=0.47), the MONS response remained positively correlated with punishment rating after controlling for the BAAS response (path $b$; β=0.37, s.e.m.=0.01, 95% CI=[0.34, 0.39]; $Z$=3.74, $P$<0.001; Cohen's $d$=0.64) and a significant mediation effect was observed (path $a \times b$; β=2.43, s.e.m.=0.14, 95% CI=[2.14, 2.69]; $Z$=3.53, $P$<0.001; Cohen's $d$=0.32). Moreover, both the total effect (path $c$; β=3.93, s.e.m.=0.24, 95% CI=[3.46, 4.40]; $Z$=3.70, $P$<0.001; Cohen's $d$=0.37) and the direct effect (path $c'$; β=1.28, s.e.m.=0.22, 95% CI=[0.85, 1.72]; $Z$=3.76, $P$<0.001; Cohen's $d$=0.13) were significant. Collectively, these findings indicate that the MONS response partially mediated the relationship between BAAS response and punishment rating. Importantly, comparable results were observed in the validation cohort (Supplementary Fig. 10a).

We subsequently constructed the reverse mediation model, in which BAAS response was specified as $M$, MONS response as $X$ and punishment rating as $Y$. Although BAAS response also showed a partial mediation effect (Supplementary Fig. 10b), the mediation strength was substantially weaker than that observed in the primary model (Supplementary Fig. 10a). Specifically, the mediation effect size in the primary model was 4.57 times larger in the discovery cohort (model 1: Cohen's *d* = 0.32; model 2: Cohen's *d* = 0.07), and a similar trend was observed in the validation cohort. Together, these findings suggest that although affective arousal contributes to punishment-related neural processing, the MONS captures variance more specifically linked to punishment tendency.

**MONS retains its predictive utility for subjective outrage after controlling for affective arousal**

Since affective arousal contributed, at least in part, to the subjective experience of outrage, as evidenced by the preceding analyses (e.g., Supplementary Table 2, Fig. 6c and Supplementary Fig. 8b), we next examined the extent to which arousal-related processes contribute to the predictive performance of the MONS. Following a previously described approach[8], BAAS responses were regressed out from the corresponding MONS responses, and the resulting adjusted MONS responses were subsequently used to discriminate different levels of subjective outrage. As shown in Supplementary Fig. 11, the MONS retained robust discriminative performance after BAAS-derived arousal-related variance was removed. In the discovery cohort (study 1, *n*=80), the adjusted MONS distinguished successive outrage intensity levels with 75%−86% accuracy (all *P*<0.001; Cohen's *d*=0.79–1.25; Supplementary Fig. 11a), and low, moderate and high outrage levels with 90%–97% accuracy (all *P*<0.001; Cohen's *d*=1.64–3.31; Supplementary Fig. 11b). These findings were largely replicated in the independent validation cohort (study 2, *n*=30), except for the level 2 versus level 1 comparison (accuracy=70%, *P*=0.09). These results indicate that although affective arousal contributes to the neural representation of outrage, the MONS retains predictive utility beyond BAAS-derived arousal-related variance, supporting its relative specificity for subjective outrage.

**Validating the whole-brain outrage signature in traditional moral judgment contexts**

Across all six datasets, the whole-brain MONS failed to reliably discriminate perceived moral wrongness (Supplementary Fig. 12 and Supplementary Table 9). Although a few contrasts reached statistical significance, these effects were either weak or occurred in the opposite direction, including the high-versus-moderate contrast in the vignette-based discovery cohort (accuracy=27%, *P*<0.001, Cohen's *d*=-0.65) and the immoral-versus-moral contrast in the first picture-based dataset (accuracy=24%, *P*=0.008, Cohen's *d*=-0.61).

### Supplementary Tables

Supplementary Table 1. Outrage, outrage consistency, arousal, moral wrongness and punishment of the 45 outrage-inducing short-form videos selected for the fMRI experiment, based on the pre-rating procedure (shown as mean±s.d.).

| Outrage | Outrage consistency | Arousal | Moral wrongness | Punishment |
|---|---|---|---|---|
| 7.35 (0.61) | 7.72 (0.45) | 7.24 (0.44) | 7.81 (0.67) | 7.44 (0.88) |

Supplementary Table 2. Stimulus-level correlations between outrage and other ratings across participants for the 45 outrage-inducing short-form videos selected for the fMRI experiment, based on the pre-rating procedure (mean±s.d.).

| | Arousal | Moral wrongness | Punishment |
|---|---|---|---|
| Outrage | 0.67 (0.13) | 0.66 (0.24) | 0.68 (0.23) |

Supplementary Table 3. Prediction–outcome correlations (mean±s.d.) with various numbers of voxels (discovery cohort).

| Number of voxels | Vis | SM | dA | vA | Limb | FP | DMN | subC | Whole-brain |
|---|---|---|---|---|---|---|---|---|---|
| 50 | 0.184 (0.040) | 0.331 (0.039) | 0.329 (0.038) | 0.322 (0.039) | 0.237 (0.038) | 0.293 (0.042) | 0.427 (0.036) | 0.283 (0.038) | 0.351 (0.045) |
| 150 | 0.234 (0.035) | 0.318 (0.035) | 0.319 (0.034) | 0.330 (0.035) | 0.251 (0.034) | 0.313 (0.039) | 0.354 (0.037) | 0.319 (0.032) | 0.358 (0.042) |
| 250 | 0.240 (0.029) | 0.303 (0.032) | 0.309 (0.030) | 0.334 (0.030) | 0.256 (0.032) | 0.315 (0.032) | 0.317 (0.033) | 0.331 (0.031) | 0.353 (0.039) |
| 500 | 0.241 (0.022) | 0.290 (0.024) | 0.292 (0.022) | 0.326 (0.024) | 0.259 (0.025) | 0.301 (0.025) | 0.293 (0.029) | 0.343 (0.027) | 0.358 (0.035) |
| 750 | 0.235* (0.018) | 0.278 (0.021) | 0.280 (0.019) | 0.315 (0.019) | 0.250* (0.020) | 0.295 (0.023) | 0.296 (0.026) | 0.347 (0.022) | 0.402 (0.032) |
| 1000 | 0.235* (0.015) | 0.269* (0.018) | 0.275* (0.017) | 0.306 (0.016) | 0.240* (0.017) | 0.292* (0.020) | 0.300 (0.023) | 0.347 (0.019) | 0.437 (0.029) |
| 2000 | 0.249* (0.011) | 0.263* (0.014) | 0.271* (0.012) | 0.289* (0.011) | 0.231* (0.012) | 0.301* (0.014) | 0.311* (0.016) | 0.353* (0.014) | 0.488 (0.023) |
| 3500 | 0.257* (0.008) | 0.270* (0.010) | 0.274* (0.008) | 0.293* (0.008) | 0.241* (0.009) | 0.310* (0.010) | 0.321* (0.011) | 0.357* (0.009) | 0.507 (0.017) |
| 6500 | 0.261* (0.005) | 0.273* (0.007) | 0.279* (0.005) | 0.297* (0.005) | 0.246* (0.005) | 0.316* (0.007) | 0.325* (0.008) | 0.354* (0.005) | 0.521 (0.013) |
| 10000 | 0.263* (0.004) | 0.274* (0.005) | 0.281* (0.003) | 0.299* (0.002) | 0.248* (0.003) | 0.319* (0.005) | 0.327* (0.006) | 0.349* (0.001) | 0.526 (0.010) |
| 14000 | NA | NA | NA | NA | NA | NA | NA | NA | 0.529 (0.009) |
| 30000 | NA | NA | NA | NA | NA | NA | NA | NA | 0.532 (0.006) |
| 65000 | NA | NA | NA | NA | NA | NA | NA | NA | 0.534 (0.004) |
| 150000 | NA | NA | NA | NA | NA | NA | NA | NA | 0.534 (0.002) |
| Full | 0.265 | 0.276 | 0.282 | 0.299 | 0.249 | 0.321 | 0.329 | 0.349 | 0.535 |

Vis, visual network; SM, somatomotor network; dA, dorsal attention network; vA, ventral attention network; Limb, limbic network; FP, frontoparietal network; DMN, default model network; subC, subcortical network; NA, not applicable. *indicates that the prediction is significantly lower as compared with the whole-brain model using the same number of voxels (two-tailed *Z*-test; $P<0.05$, Bonferroni corrected).

Supplementary Table 4. Prediction–outcome correlations (mean±s.d.) with various numbers of voxels (validation cohort).

| Number of voxels | Vis | SM | dA | vA | Limb | FP | DMN | subC | Whole-brain |
|---|---|---|---|---|---|---|---|---|---|
| 50 | 0.177 (0.052) | 0.316 (0.046) | 0.332 (0.049) | 0.325 (0.055) | 0.312 (0.052) | 0.332 (0.052) | 0.391 (0.040) | 0.326 (0.040) | 0.363 (0.054) |
| 150 | 0.205 (0.045) | 0.311 (0.042) | 0.319 (0.045) | 0.317 (0.046) | 0.320 (0.048) | 0.351 (0.049) | 0.344 (0.047) | 0.342 (0.033) | 0.359 (0.054) |
| 250 | 0.199 (0.040) | 0.301 (0.040) | 0.301 (0.042) | 0.318 (0.044) | 0.300 (0.046) | 0.355 (0.046) | 0.310 (0.047) | 0.333 (0.032) | 0.351 (0.052) |
| 500 | 0.179 (0.034) | 0.297 (0.036) | 0.271 (0.036) | 0.296 (0.035) | 0.267 (0.039) | 0.323 (0.040) | 0.276 (0.041) | 0.315 (0.025) | 0.337 (0.050) |
| 750 | 0.172* (0.029) | 0.285 (0.032) | 0.252 (0.032) | 0.282 (0.030) | 0.255 (0.035) | 0.295 (0.036) | 0.262 (0.036) | 0.302 (0.022) | 0.365 (0.047) |
| 1000 | 0.160* (0.028) | 0.273 (0.029) | 0.246 (0.029) | 0.265 (0.028) | 0.249 (0.033) | 0.280 (0.032) | 0.261 (0.031) | 0.289 (0.020) | 0.394 (0.043) |
| 2000 | 0.154* (0.020) | 0.274* (0.021) | 0.243* (0.021) | 0.218* (0.020) | 0.245* (0.027) | 0.269* (0.023) | 0.281* (0.024) | 0.261* (0.014) | 0.436 (0.032) |
| 3500 | 0.159* (0.014) | 0.284* (0.016) | 0.244* (0.016) | 0.204* (0.013) | 0.245* (0.020) | 0.277* (0.016) | 0.290* (0.016) | 0.255* (0.010) | 0.453 (0.026) |
| 6500 | 0.161* (0.010) | 0.289* (0.010) | 0.250* (0.010) | 0.209* (0.008) | 0.262* (0.012) | 0.284* (0.011) | 0.295* (0.011) | 0.252* (0.005) | 0.463 (0.019) |
| 10000 | 0.163* (0.007) | 0.291* (0.007) | 0.251* (0.007) | 0.211* (0.003) | 0.265* (0.007) | 0.288* (0.007) | 0.297* (0.008) | 0.248* (0.001) | 0.467 (0.015) |
| 14000 | NA | NA | NA | NA | NA | NA | NA | NA | 0.470 (0.013) |
| 30000 | NA | NA | NA | NA | NA | NA | NA | NA | 0.472 (0.009) |
| 65000 | NA | NA | NA | NA | NA | NA | NA | NA | 0.473 (0.005) |
| 150000 | NA | NA | NA | NA | NA | NA | NA | NA | 0.474 (0.002) |
| Full | 0.165 | 0.293 | 0.252 | 0.211 | 0.265 | 0.291 | 0.300 | 0.248 | 0.474 |

Vis, visual network; SM, somatomotor network; dA, dorsal attention network; vA, ventral attention network; Limb, limbic network; FP, frontoparietal network; DMN, default model network; subC, subcortical network; NA, not applicable. *indicates that the prediction is significantly lower as compared with the whole-brain model using the same number of voxels (two-tailed *Z*-test; $P<0.05$, Bonferroni corrected).

Supplementary Table 5. Information for each study.

| *Study* | *Dataset ref.* | *Stimuli Types* | *N (female)* | *Mean age (s.d.)* | *Study location* | *MRI system* |
|---|---|---|---|---|---|---|
| 1 | Discovery[a] | Videos with varying outrage intensity levels | 80 (40) | 21.89 (2.45) | UESTC | 3T GE |
| 2 | Validation[a] | Videos with varying outrage intensity levels | 30 (16) | 20.80 (2.67) | UESTC | 3T GE |
| 3 | Wagels et al.[1] and Akkoc Altinok et al.[2] | Aggression following high versus low provocation | Group1[b]: 41 (0)<br>Group2[c]: 27 (0) | Group1: 23.74 (3.48)<br>Group2: 24.46 (3.81) | Aachen | 3T Siemens Prisma |
| 4 | Gan et al.[3] | Visual disgust stimuli with varying disgust intensity levels | 78 (44) | 22.10 (2.68) | UESTC | 3T GE |
| 5 | Gan et al.[3] | Visual disgust stimuli with varying disgust intensity levels | 30 (16) | 21.13 (2.18) | UESTC | 3T GE |
| 6 | Čeko et al.[7] | Pressure pain with intensity levels<br>Thermal pain with intensity levels<br>Aversive sounds with intensity levels<br>Aversive images with intensity levels | 55 (24) | 24.9 (6.8) | University of Colorado Boulder | 3T Siemens Prisma |
| 7 | Zhang et al.[8] | Videos with varying arousal intensity levels | 36 (17) | 21.03 (2.22) | UESTC | 3T GE |
| 8 | Hopp et al.[9] (Discovery) | Moral vignettes with varying wrongness levels | 64 (31) | 20.78 (2.45) | UCSB | 3T Siemens Prisma |
| 9 | Hopp et al.[9] (Validation) | Moral vignettes with varying | 30 (19) | 20.51 (2.35) | UCSB | 3T Siemens Prisma |

| | | | | | | |
|---|---|---|---|---|---|---|
| | | wrongness levels | | | | |
| 10 | Hopp et al.[9] (Replication) | Moral vignettes with varying wrongness levels | 27 (13) | 24.65 (4.21) | Duke | 3T GE |
| 11 | Hopp et al.[9] (Generalization) | Moral vignettes with varying wrongness levels | 30 (18) | 26.4 (9.17) | University of Amsterdam | 3T Philips Achiva |
| 12 | Hopp et al.[9] (Picture dataset 1) | Moral and immoral pictures | 30 (19) | 20.51 (2.35) | UCSB | 3T Siemens Prisma |
| 13 | Hopp et al.[9] (Picture dataset 2) | Moral and immoral pictures | 30 (18) | 26.4 (9.17) | University of Amsterdam | 3T Philips Achiva |

[a] current study data; [b] post-task state anger increased; [c] post-task state anger did not increase.

Supplementary Table 6. Prediction (correlation) of MONS in the disgust context.

| Datasets | Overall (bootstrapped 95% CI) and within-subject (mean±s.e.m.) prediction–outcome *r* |
|---|---|
| Study 4: Disgust discovery cohort | 0.21 [0.11, 0.30]; 0.44±0.06 |
| Study 5: Disgust validation cohort | 0.19 [0.02, 0.34]; 0.44±0.08 |

We applied the MONS to two disgust datasets (i.e., studies 4,5) and calculated the overall (bootstrapped 95% CI) as well as within-subject (mean±s.e.m.) prediction–outcome correlations between the pattern expressions and the true ratings.

Supplementary Table 7. Selection of ROIs for Fig. 6a,b.

| ROI | Areas included in ROI | Atlas |
|---|---|---|
| *Cortical* | | |
| Dorsolateral prefrontal cortex (dLPFC) | 8Av, p9-46v, 46 | HCP-MMP1.0, Glasser et al. 2016[15] |
| Ventrolateral prefrontal cortex (vLPFC) | 44, 45 | HCP-MMP1.0, Glasser et al. 2016[15] |
| Dorsomedial prefrontal cortex (dMPFC) | 8BM, 8BL, 9m | HCP-MMP1.0, Glasser et al. 2016[15] |
| Ventromedial prefrontal cortex (vMPFC) | 10r, 10v, 10d, 10pp, OFC | HCP-MMP1.0, Glasser et al. 2016[15] |
| Orbitofrontal cortex (OFC) | 11l, 47s/47m, 47l, a47r, p47r | HCP-MMP1.0, Glasser et al. 2016[15] |
| Anterior cingulate cortex (ACC) | Cingulum_Ant_L, Cingulum_Ant_R | AAL, Tzourio-Mazoyer et al., 2002[24] |
| Middle cingulate cortex (MCC) | Cingulum_Mid_L, Cingulum_Mid_R | AAL, Tzourio-Mazoyer et al., 2002[24] |
| Posterior cingulate cortex (PCC) | Cingulum_Post_L, Cingulum_Post_R | AAL, Tzourio-Mazoyer et al., 2002[24] |
| Anterior insula (aINS) | FOP4, FOP5, AVI, AAIC | HCP-MMP1.0, Glasser et al. 2016[15] |
| Middle insula (mINS) | MI, PoI2, FOP2, FOP3 | HCP-MMP1.0, Glasser et al. 2016[15] |
| Temporoparietal junction (TPJ) | Temporoparietal Junction | Mars et al., 2012[25] |
| Precuneus | PCun_L(R)_4_1, PCun_L(R)_4_2, PCun_L(R)_4_3, PCun_L(R)_4_4 | Human Brainnetome Atlas, Fan et al., 2016[26] |
| *Subcortical* | | |
| Periaqueductal gray (PAG) | PAG | Shen et al., 2013[27] |

| | | |
|---|---|---|
| Hypothalamus | Hythal | HCP-MMP1.0, Glasser et al. 2016[15] |
| Amygdala | Centro-medial, latero-basal, superficial | SPM Anatomy Toolbox v2.2c, Eickhoff et al.2005[28] |
| Thalamus | Tha_L(R)_8_1, Tha_L(R)_8_2, Tha_L(R)_8_3, Tha_L(R)_8_4, Tha_L(R)_8_5, Tha_L(R)_8_6, Tha_L(R)_8_7, Tha_L(R)_8_8 | Human Brainnetome Atlas, Fan et al., 2016[26] |
| Ventral striatum (vStr) | BG_L(R)_6_3 | Human Brainnetome Atlas, Fan et al., 2016[26] |

Supplementary Table 8. Prediction (correlation) of MONS without visual and auditory cortices in the context of moral vignettes.

| Datasets | Overall (bootstrapped 95% CI) and within-subject (mean±s.e.m.) prediction–outcome *r* |
|---|---|
| Study 8: Moral vignettes discovery cohort | 0.09 [-0.03, 0.21]; 0.15±0.08 |
| Study 9: Moral vignettes validation cohort | 0.06 [-0.12, 0.24]; 0.20±0.11 |
| Study 10: Moral vignettes replication cohort | 0.01 [-0.19, 0.22]; 0.05±0.11 |
| Study 11: Moral vignettes generalization cohort | 0.03 [-0.15, 0.21]; 0.09±0.12 |

We applied the MONS without visual and auditory cortices to four moral vignette datasets (i.e., studies 8 to 11) and calculated the overall (bootstrapped 95% CI) as well as within-subject (mean±s.e.m.) prediction–outcome correlations between the pattern expressions and the true ratings.

Supplementary Table 9. Prediction (correlation) of MONS in the context of moral vignettes.

| Datasets | Overall (bootstrapped 95% CI) and within-subject (mean±s.e.m.) prediction−outcome *r* |
|---|---|
| Study 8: Moral vignettes discovery cohort | -0.04 [-0.16, 0.09]; -0.09±0.08 |
| Study 9: Moral vignettes validation cohort | -0.002 [-0.18, 0.18]; -0.006±0.11 |
| Study 10: Moral vignettes replication cohort | -0.03 [-0.23, 0.17]; -0.09±0.11 |
| Study 11: Moral vignettes generalization cohort | -0.02 [-0.20, 0.17]; 0.003±0.12 |

We applied the MONS to four moral vignette datasets (i.e., studies 8 to 11) and calculated the overall (bootstrapped 95% CI) as well as within-subject (mean±s.e.m.) prediction–outcome correlations between the pattern expressions and the true ratings.

### Supplementary Figures

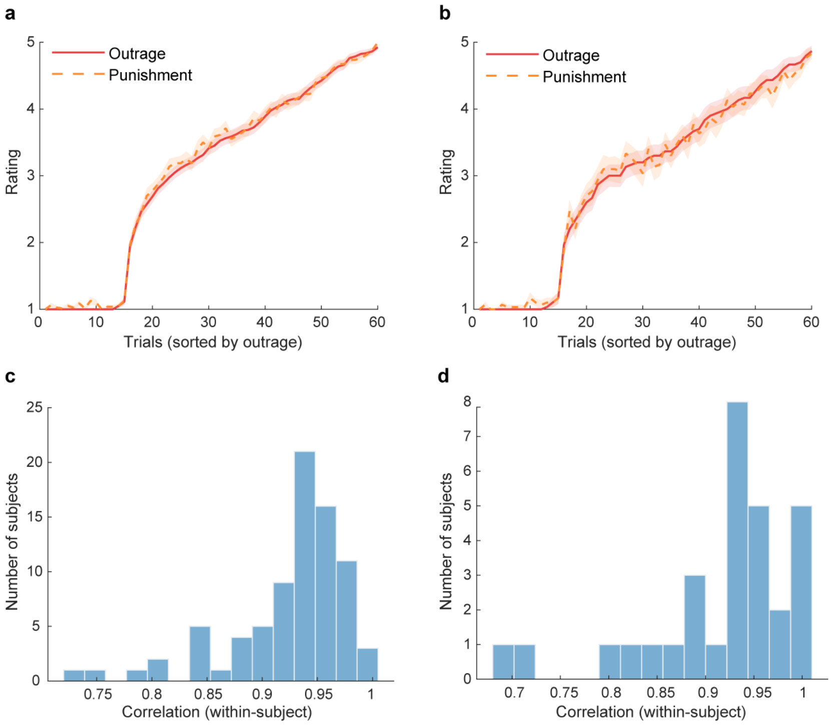

**Supplementary Fig. 1 | Aligned trial-wise profiles of outrage and punishment ratings in discovery and validation cohorts. a**,**b**, Trial-wise profiles of outrage (red) and punishment (orange) ratings after sorting trials in ascending order of outrage ratings within each participant in the discovery cohort (**a**; *n*=80) and independent validation cohort (**b**; *n*=30), respectively. Shaded areas represent s.e.m. This sorting procedure aligns trials according to subjective outrage intensity, enabling a direct comparison of the corresponding punishment ratings. The close overlap between the two curves in both cohorts indicates that punishment ratings closely track outrage ratings across trials, suggesting that the two measures share a highly similar trial-wise structure. **c**,**d**, Distribution of within-subject correlations between the two measures in the discovery (**c**) and validation (**d**) cohorts, respectively.

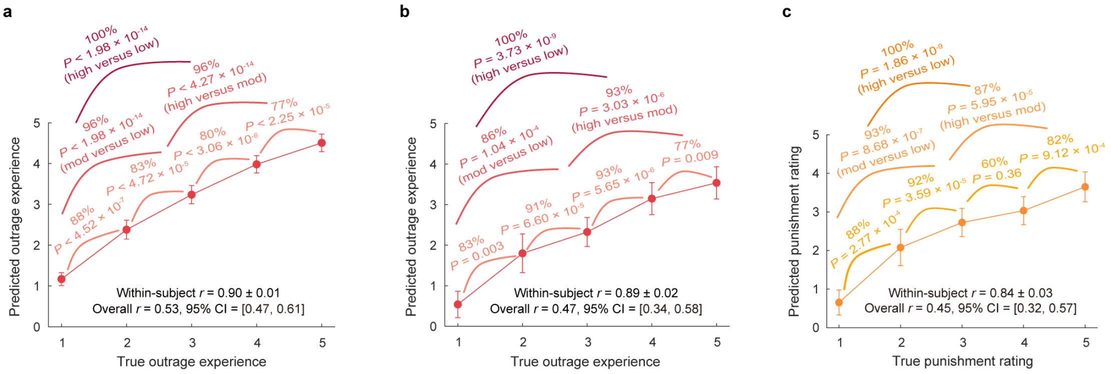

**Supplementary Fig. 2 | MONS prediction after excluding visual and auditory cortices. a**, Predicted versus actual outrage ratings in the cross-validated discovery cohort (*n*=80). **b**, Predicted versus actual outrage ratings in the independent validation cohort (*n*=30). **c**, Predicted versus actual punishment ratings in the independent validation cohort (*n*=30). Classification accuracies reflect forced-choice comparisons across different rating levels. *P* values are based on two-sided binomial tests. *r* indicates the Pearson correlation coefficient between predicted and actual ratings. Error bars reflect the s.e.m.

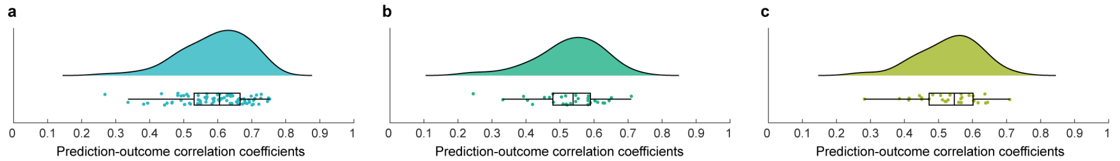

**Supplementary Fig. 3 | Within-subject trial-wise prediction using original ratings. a**, Distribution of within-subject trial-wise Pearson correlations between predicted and actual outrage ratings across participants in the discovery cohort; significant prediction–outcome correlations were observed in 79 of 80 participants ($P$=[1.75×$10^{-12}$, 0.047]; $P$>0.05 for the remaining participant). **b**, Distribution of within-subject trial-wise Pearson correlations between predicted and actual outrage ratings across participants in the validation cohort; significant prediction–outcome correlations were observed in 29 of 30 participants ($P$=[6.43×$10^{-10}$, 0.014]; $P$=0.07 for the remaining participant). **c**, Distribution of within-subject trial-wise Pearson correlation between predicted and actual punishment ratings across participants in the validation cohort; significant prediction–outcome correlations were observed in all 30 participants ($P$=[6.54×$10^{-10}$, 0.033]). **a**–**c**, In the raincloud plots, boxplots indicate the interquartile range (IQR), with the center line representing the median, box boundaries indicating the 25th and 75th percentiles, and the whiskers extending to the minimum and maximum values within 1.5 × IQR from the quartiles.

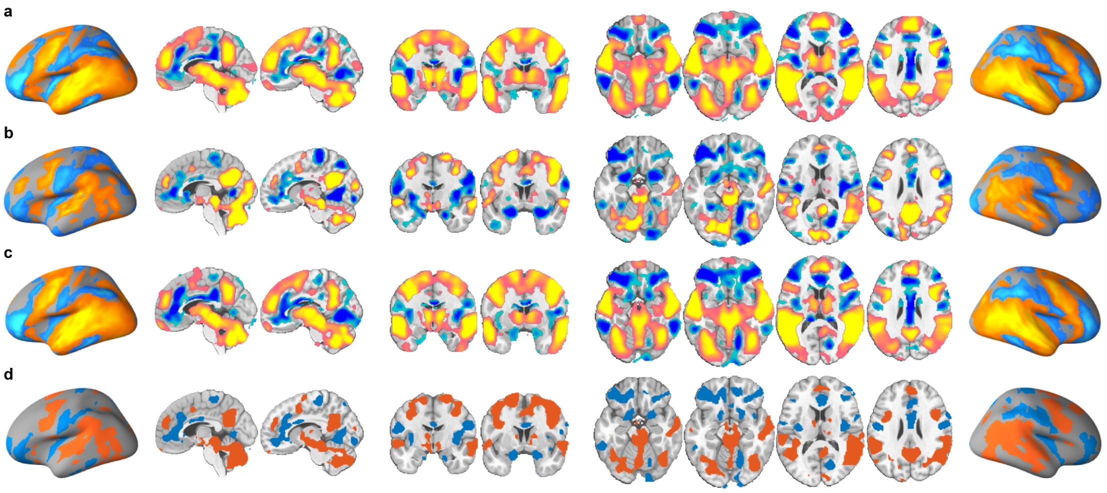

**Supplementary Fig. 4** | Subjective experience of outrage is associated with and predicted by distributed brain regions. **a**, Univariate parametric effects of outrage ratings. **b**, Within-subject outrage-predictive patterns. **c**, Thresholded 'activation patterns' reconstructed from within-subject outrage-predictive patterns. **d**, Conjunction between the predictive-weight patterns and reconstructed activation patterns shown in (**b**) and (**c**). Images thresholded at $q<0.05$, FDR corrected. Hot color indicates positive associations (**a** and **c**) or predictive weights (**b**), whereas cold color indicates negative associations (**a** and **c**) or predictive weights (**b**).

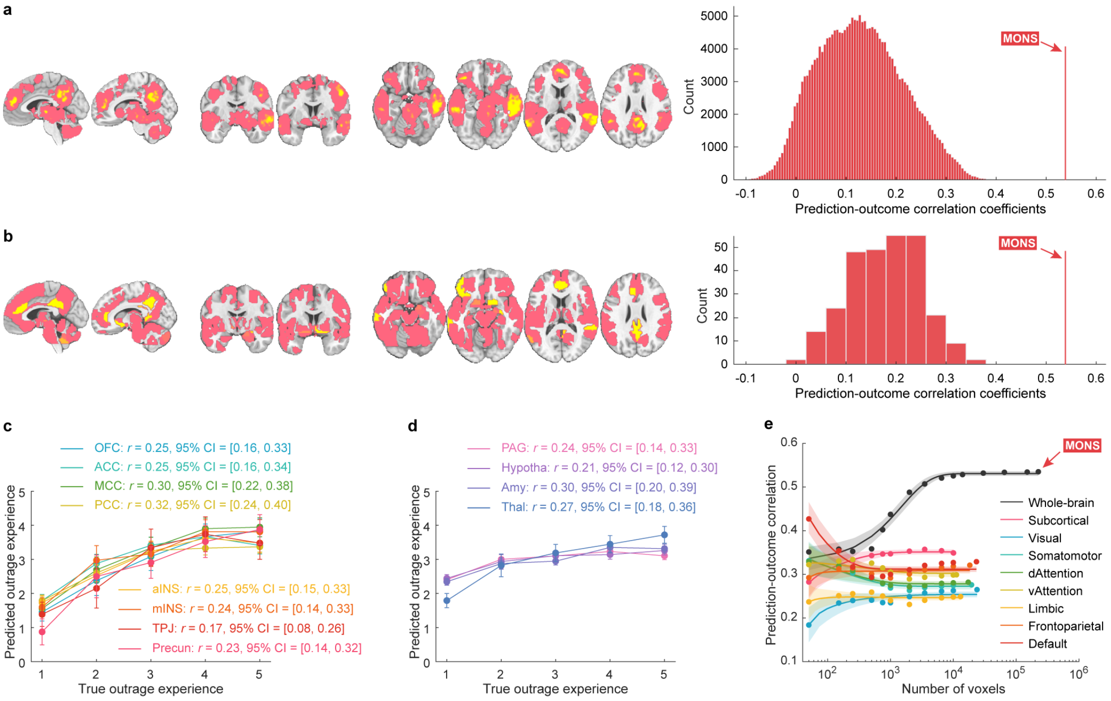

**Supplementary Fig. 5 | Prediction of subjective outrage from local brain regions and large-scale networks in the discovery cohort. a,b**, Brain regions showing significant prediction of subjective outrage identified using searchlight- and parcellation-based analyses, respectively. Statistical significance was assessed using two-sided Pearson prediction–outcome correlation. Because uncorrected *P* values corresponding to *q*<0.05 may be relatively lenient, only regions surviving *P*<0.001 uncorrected (corresponding to FDR-corrected *q*<0.004 and *q*<0.002 for the searchlight- and parcellation-based predictions, respectively) are displayed. Histograms depict the distribution of cross-validated predictions (correlations) across local searchlights or atlas-defined parcellations. Red lines indicate the prediction–outcome correlation from the whole-brain MONS. **c,d**, Cross-validated predictions (mean±s.e.m.) obtained from cortical ROI- and subcortical ROI-based prediction analyses, respectively. Error bar represents s.e.m.; *r* denotes the overall (between- and within-subjects) prediction–outcome Pearson correlation coefficient. **e**, Subjective experience of outrage is represented across distributed systems. Model performance was evaluated as increasing numbers of voxels/features (*x*-axis) were randomly sampled to predict subjective outrage from the whole-brain (black), subcortical network (pink), or individual large-scale cerebral networks (other colored lines). The *y*-axis indicates the cross-validated prediction–outcome correlation. Colored dots denote mean correlation coefficients, solid lines indicate the mean parametric fit, and shaded areas represent the s.d.

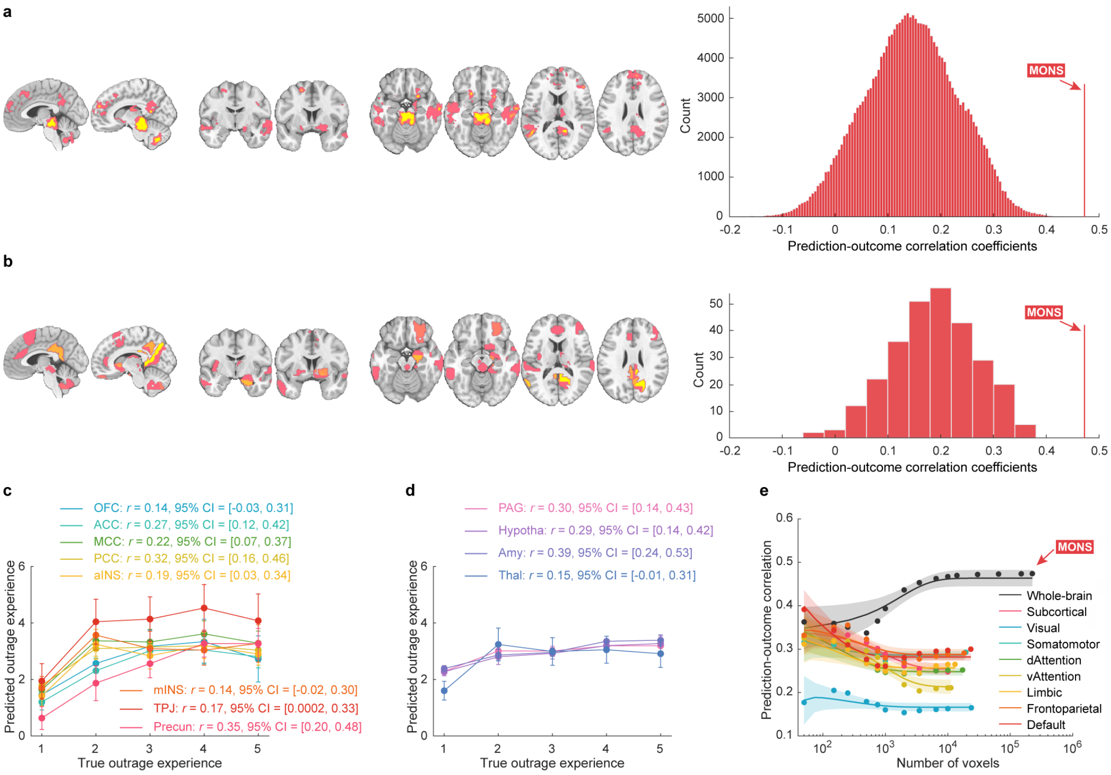

**Supplementary Fig. 6 | Predictions of models trained on the discovery cohort on the validation cohort. a,b**, Brain regions showing significant prediction of subjective outrage identified using searchlight- and parcellation-based analyses, respectively. Statistical significance was evaluated by two-sided Pearson prediction–outcome correlation. Only regions surviving *P*<0.001 uncorrected are displayed. Histograms: Predictions (correlations) from searchlights (Panel a) and parcellations (Panel b), respectively. Red lines indicate the prediction–outcome correlation from the MONS. **c,d**, Predictions (mean±s.e.m.) from cortical ROI- and subcortical ROI-based prediction analyses, respectively. Error bar represents the s.e.m.; *r* indicates the overall (between- and within-subjects) prediction–outcome Pearson correlation coefficient. **e**, Subjective outrage is distributed across multiple systems. Model performance was assessed as increasing numbers of voxels/features (*x*-axis) were randomly sampled to predict subjective outrage from the whole brain (black), subcortical network (pink), or individual large-scale cerebral networks (other colored lines). The *y*-axis denotes the prediction–outcome correlation. Colored dots represent mean correlation coefficients, solid lines indicate the mean parametric fit and shaded areas represent the s.d.

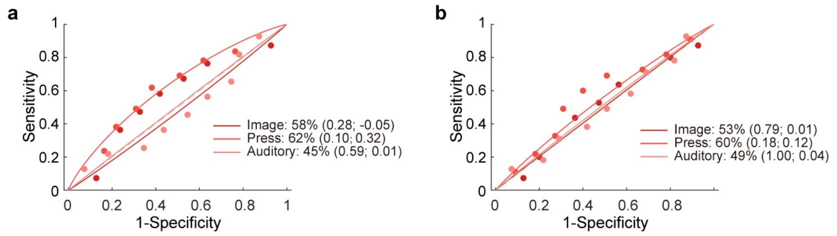

**Supplementary Fig. 7 | Classification performance of the MONS in additional multimodal negative affect contexts. a**, The MONS does not predict high versus low aversive image, mechanical pain, and aversive sound conditions, respectively (shown as forced-choice classification accuracy, *P* value and Cohen's *d*). **b**, The MONS does not predict high versus moderate aversive image, mechanical pain, and aversive sound conditions, respectively. *P* values were derived from binomial tests, two-sided.

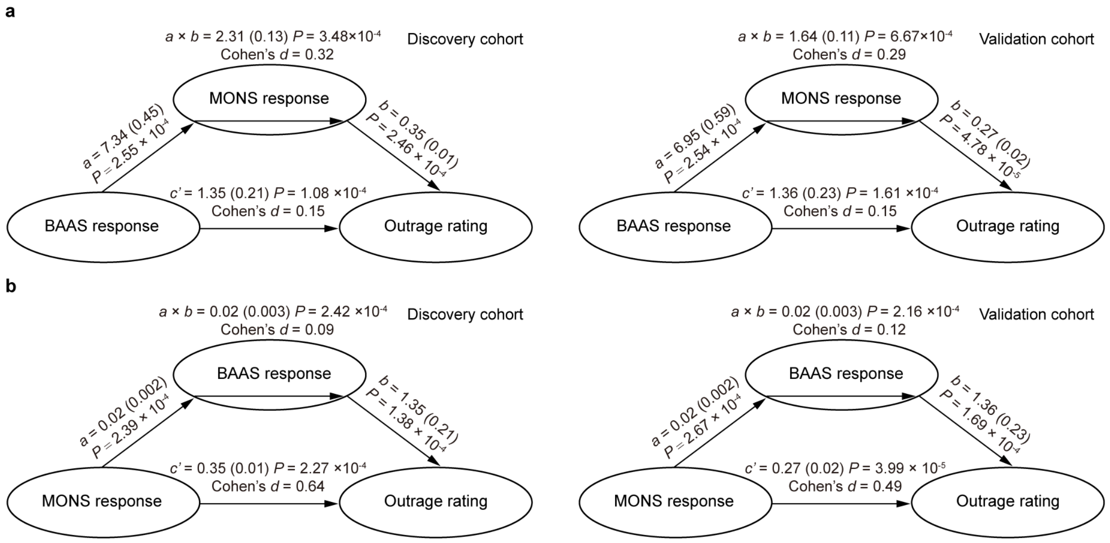

**Supplementary Fig. 8 | Multilevel mediation analyses exploring the relationship between MONS response, BAAS response and subjective outrage rating. a**, MONS response partially mediates the effect of BAAS response on the subjective outrage rating. **b**, BAAS response partially mediates the association between MONS response and subjective outrage rating. **a,b**, Two-sided *P* values were derived from bootstrap tests with 10,000 samples.

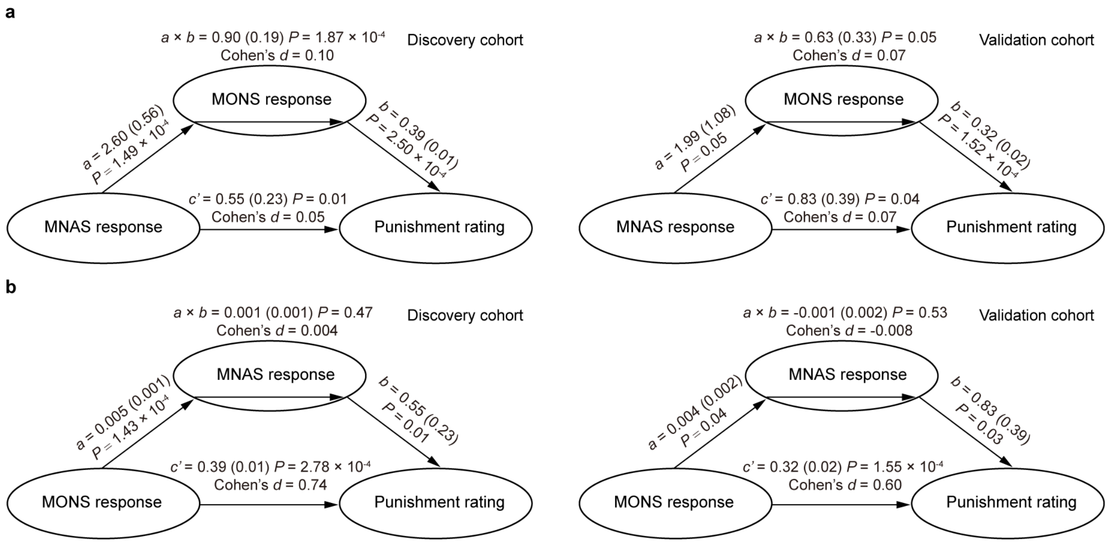

**Supplementary Fig. 9 | Multilevel mediation analyses exploring the relationship between MONS response, MNAS response and punishment rating. a**, MONS response partially mediates the effect of MNAS response on punishment rating. **b**, MNAS response does not mediate the association between MONS response and punishment rating. **a,b**, Two-sided *P* values were estimated using bootstrap tests with 10,000 samples.

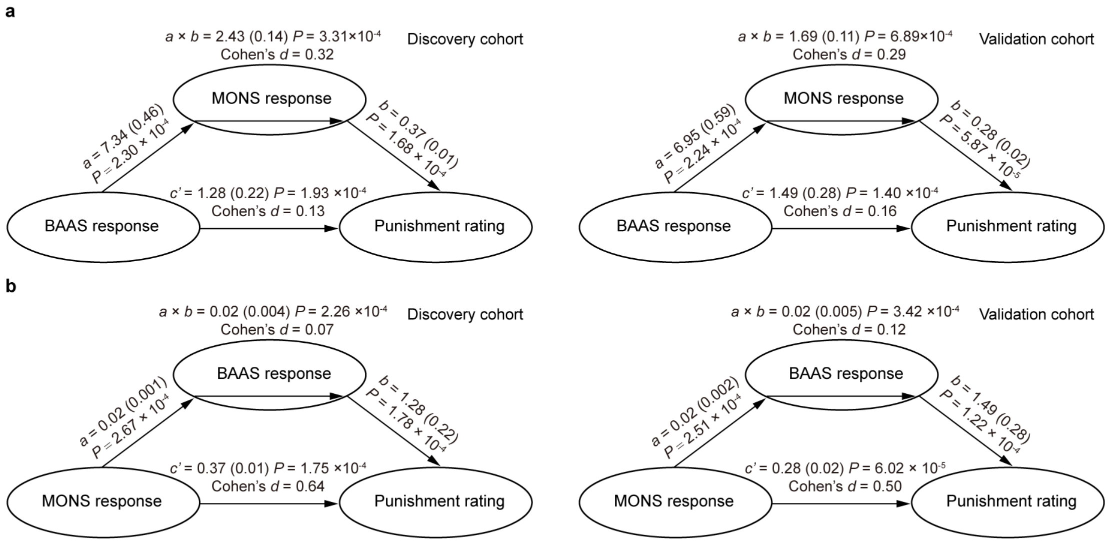

**Supplementary Fig. 10 | Multilevel mediation analyses exploring the relationship between MONS response, BAAS response and punishment rating. a**, MONS response partially mediates the effect of BAAS response on punishment rating. **b**, BAAS response partially mediates the association between MONS response and punishment rating. **a,b**, Two-sided *P* values were derived from bootstrap tests with 10,000 samples.

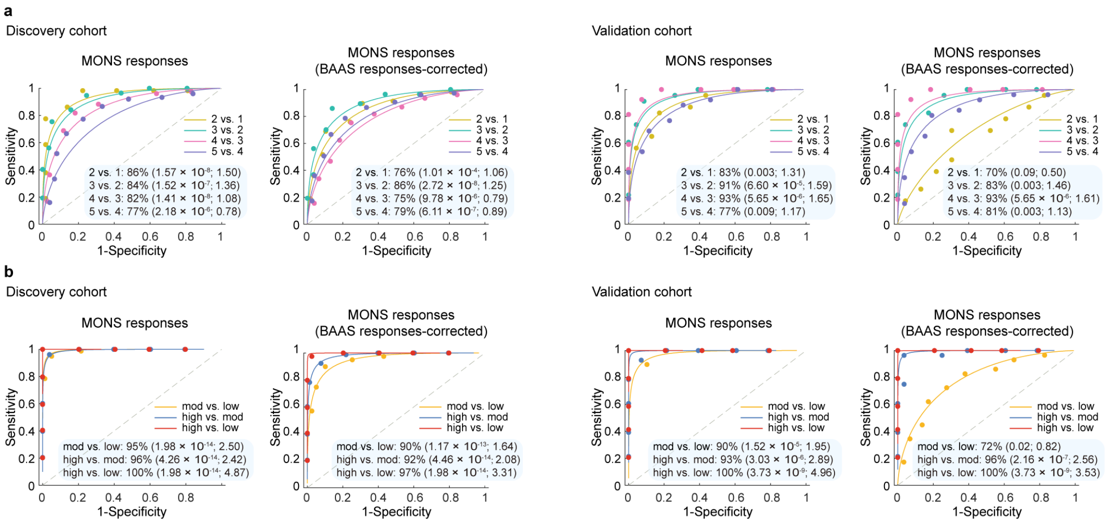

**Supplementary Fig. 11 | Evaluating the specificity of MONS after controlling for affective arousal.** **a**, MONS retains robust discriminative performance across successive outrage intensity levels after BAAS-response corrected in both the discovery and validation cohorts (shown as forced-choice classification accuracy, *P* value and Cohen's *d*), except for the level 2 versus level 1 comparison in the validation cohort. **b**, MONS retains robust discriminative performance across low, moderate and high outrage intensity levels after BAAS-response corrected in both the discovery and validation cohorts. *P* values were based on binomial tests, two-sided.

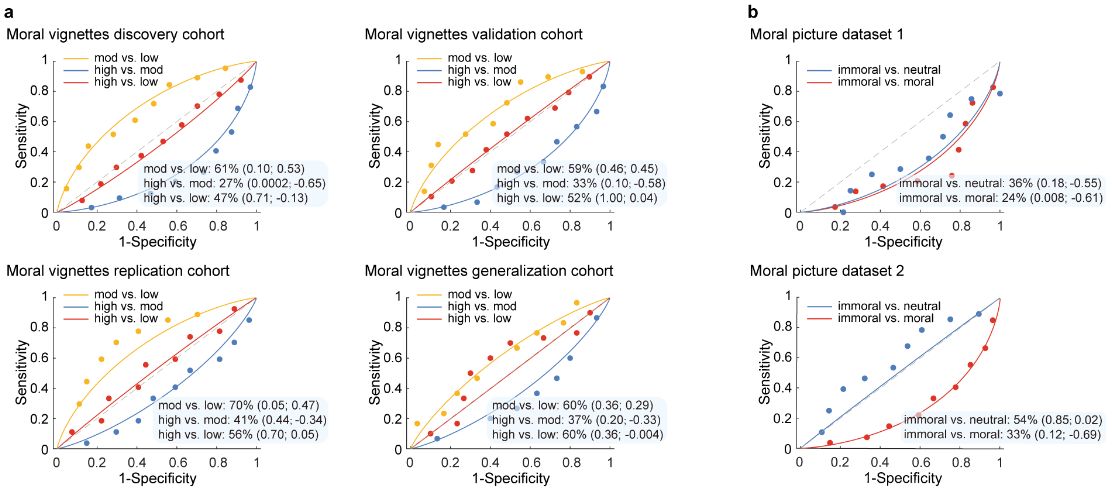

**Supplementary Fig. 12 | Validation of the whole-brain MONS in traditional moral judgment contexts. a**, The MONS failed to reliably distinguish different levels of moral wrongness across the four vignette-based moral judgment datasets, except for the comparison between moderate and low moral wrongness in the moral vignette-based replication cohort, which reached marginal significance (*P*=0.05) (shown as forced-choice classification accuracy, *P* value and Cohen's *d*). **b**, The MONS inaccurately discriminated between immoral and moral or neutral conditions in two picture-based moral judgment datasets. Note that significant predictions observed for high versus moderate moral wrongness in the moral vignette-based discovery cohort and for immoral versus moral classification in the first moral picture dataset occurred in the opposite direction. *P* values were based on binomial tests, two-sided.

### Supplemental references

1. Wagels, L., et al. Exogenous testosterone and the monoamine-oxidase A polymorphism influence anger, aggression and neural responses to provocation in males. *Neuropharmacology* **156**, 107491 (2019).
2. Akkoc Altinok, D. C., et al. A combined administration of testosterone and arginine vasopressin affects aggressive behavior in males. *Brain Sci.* **11**, 1623 (2021).
3. Gan, X., et al. A neurofunctional signature of subjective disgust generalizes to oral distaste and socio-moral contexts. *Nat. Hum. Behav.* **8**, 1383-1402 (2024).
4. Haberkamp, A., Glombiewski, J. A., Schmidt, F. & Barke, A. The DIsgust-RelaTed-Images (DIRTI) database: validation of a novel standardized set of disgust pictures. *Behav. Res. Ther.* **89**, 86-94 (2017).
5. Marchewka, A., Żurawski, Ł., Jednoróg, K. & Grabowska, A. The Nencki Affective Picture System (NAPS): introduction to a novel, standardized, wide-range, high-quality, realistic picture database. *Beh. Res. Meth.* **46**, 596-610 (2014).
6. Lang, P., Bradley, M. & Cuthbert, B. *International Affective Picture System (IAPS): affective ratings of pictures and instruction manual*. Technical Report A-8. University of Florida, Gainesville (2008).
7. Čeko, M., Kragel, P. A., Woo, C.-W., López-Solà, M. & Wager, T. D. Common and stimulus-type-specific brain representations of negative affect. *Nat. Neurosci.* **25**, 760-770 (2022).
8. Zhang, R., et al. A neurofunctional signature of affective arousal generalizes across valence domains and distinguishes subjective experience from autonomic reactivity. *Nat. Commun.* **16**, 6492 (2025).
9. Hopp, F., Youk, S., Sinnott-Armstrong, W. & Weber, R. A sensitive and specific neural signature robustly predicts graded computations of moral wrongness. *Prepint*, (2025).
10. Clifford, S., Iyengar, V., Cabeza, R. & Sinnott-Armstrong, W. Moral foundations vignettes: a standardized stimulus database of scenarios based on moral foundations theory. *Beh. Res. Meth.* **47**, 1178-1198 (2015).
11. Hopp, F. R., Jargow, B., Kouwen, E. & Bakker, B. N. The Dutch moral foundations stimulus database: An adaptation and validation of moral vignettes and sociomoral images in a Dutch sample. *Judgment and Decision Making* **19**, e10 (2024).
12. Curry, O. S., Jones Chesters, M. & Van Lissa, C. J. Mapping morality with a compass: Testing the theory of 'morality-as-cooperation' with a new questionnaire. *Journal of Research in Personality* **78**, 106-124 (2019).
13. Crone, D. L., Bode, S., Murawski, C. & Laham, S. M. The Socio-Moral Image Database (SMID): A novel stimulus set for the study of social, moral and affective processes. *PLoS ONE* **13**, e0190954 (2018).
14. Koban, L., Jepma, M., López-Solà, M. & Wager, T. D. Different brain networks mediate the effects of social and conditioned expectations on pain. *Nat. Commun.* **10**, 4096 (2019).
15. Glasser, M. F., et al. A multi-modal parcellation of human cerebral cortex. *Nature* **536**, 171-178 (2016).
16. Haufe, S., et al. On the interpretation of weight vectors of linear models in multivariate neuroimaging. *Neuroimage* **87**, 96-110 (2014).
17. Alia-Klein, N., et al. The feeling of anger: from brain networks to linguistic expressions. *Neurosci. Biobehav. Rev.* **108**, 480-497 (2020).

18. Blair, R. J., Habel, U., Votinov, M., Wagels, L. & Denson, T. F. A cognitive neuroscience approach to understanding aggression and its treatment. *Neurosci. Biobehav. Rev.* **179**, 106386 (2025).
19. Gan, X., et al. Does unfairness evoke anger or disgust? A quantitative neurofunctional dissection based on 25 years of neuroimaging. *Neurosci. Biobehav. Rev.* **178**, 106356 (2025).
20. Klaus, J. & Schutter, D. J. L. G. Functional topography of anger and aggression in the human cerebellum. *Neuroimage* **226**, 117582 (2021).
21. Satpute, A. B. & Lindquist, K. A. The default mode network's role in discrete emotion. *Trends Cogn. Sci.* **23**, 851-864 (2019).
22. Zhou, F., et al. A distributed fMRI-based signature for the subjective experience of fear. *Nat. Commun.* **12**, 6643 (2021).
23. Kragel, P. A., Koban, L., Barrett, L. F. & Wager, T. D. Representation, pattern information, and brain signatures: from neurons to neuroimaging. *Neuron* **99**, 257-273 (2018).
24. Tzourio-Mazoyer, N., et al. Automated Anatomical Labeling of Activations in SPM Using a Macroscopic Anatomical Parcellation of the MNI MRI Single-Subject Brain. *Neuroimage* **15**, 273-289 (2002).
25. Mars, R. B., Sallet, J., Schüffelgen, U., Jbabdi, S., Toni, I. & Rushworth, M. F. Connectivity-based subdivisions of the human right "temporoparietal junction area": evidence for different areas participating in different cortical networks. *Cereb. Cortex* **22**, 1894-1903 (2012).
26. Fan, L., et al. The human brainnetome atlas: a new brain atlas based on connectional architecture. *Cereb. Cortex* **26**, 3508-3526 (2016).
27. Shen, X., Tokoglu, F., Papademetris, X. & Constable, R. T. Groupwise whole-brain parcellation from resting-state fMRI data for network node identification. *Neuroimage* **82**, 403-415 (2013).
28. Eickhoff, S. B., et al. A new SPM toolbox for combining probabilistic cytoarchitectonic maps and functional imaging data. *Neuroimage* **25**, 1325-1335 (2005).
